# Calcium homeostasis disruption initiates rapid growth after micro-fragmentation in the scleractinian coral *Porites lobata*

**DOI:** 10.1101/2022.02.28.482414

**Authors:** Colin Lock, Bastian Bentlage, Laurie J Raymundo

**Affiliations:** Marine Laboratory, University of Guam, Mangilao, GU 96913, USA

**Keywords:** transcriptome, gene expression, calcium homeostasis, stress response, coral

## Abstract

Coral reefs are ecosystems under increasing threat from global climate change. Coral restoration is a tool for preserving biological and ecological function of coral reefs by mitigating coral loss and maintaining the structural integrity and complexity of reefs. To generate the necessary stock for coral restoration, larger coral colonies are usually fragmented to generate smaller specimens for outplanting, taking advantage of the high regenerative ability of corals. In this study, we utilized RNA-seq technology to understand the physiological responses of *Porites lobata* colonies to physical fragmentation and outplanting, which have thus far not been characterized. Our results demonstrate that *P. lobata* fragments undergoing physical injury recover through two distinct phases: rapid wound regeneration of the cut margins, followed by a slower growth phase that cements the colony to the substrate. Our study found *rapid* physiological responses to acute physical injury and outplanting in the coral host that involved significantly increased energy production, calcium homeostasis disruption, and Endoplasmic Reticulum (ER) stress leading to increased antioxidant expression and rates of protein turnover. Our results suggest that phosphoinositide-mediated acute calcium homeostasis disruption stimulates wound recovery processes in response to physical injury. Symbiont gene expression revealed extremely low gene differences in response to fragmentation, growth, and outplanting. These results provide insight into the physiological mechanisms that allow for rapid wound healing and stabilization in response to physical injury in corals.

## 1 INTRODUCTION

Global coral reef decline has become so pronounced that passive conservation strategies that rely on natural recovery of reefs, such as the designation of marine protected areas, may be inadequate to preserve the biodiversity and ecosystem services tropical coral reefs provide (Rinkevich, 2005; Forsman et al., 2006). To address this concern, active coral restoration is rapidly becoming an integral management tool to mitigate coral loss and maintain the structural integrity and complexity of reefs (Kojis & Quinn, 2001; Baums, 2008; Edwards, 2008; Barton et al., 2017). Procuring fragments from healthy coral colonies and propagating these to establish coral nursery stock for outplanting in restoration efforts is a commonly used approach in coral reef restoration (Epstein et al., 2001; Rinkevich, 2005; Edwards & Gomez, 2011). After nursery stock has reached a stable size and thus an increased chance of long-term survival (Lirman, 2000; Rinkevich, 2005; Edwards & Gomez, 2011), fragments may be cut from the coral stock.

These fragments are then firmly affixed to degraded reefs, while sustaining the initial broodstock for future outplanting (Epstein et al., 2001). While fragmentation has been successfully used as a coral reef restoration technique for corals that fragment naturally, such as species of *Acropora*, (Lirman, 2000; Highsmith, 2007; Lirman et al., 2010), micro-fragmentation of corals that do not naturally propagate asexually via fragmentation, such as species of *Porites*, has been a more recent development (Forsman et al., 2006, 2015; Page et al., 2018). Micro-fragmentation involves cutting corals into minute pieces and growing them in a protected nursery, during which corals grow to a size suitable for outplanting (Forsman et al., 2006, 2015; Page et al., 2018). Micro-fragmentation allows the generation of a large number of fragments for stress experiments or outplanting, while minimizing impacts to source colonies.

Acute physical injury, such as that caused by corallivorous fish and fragmentation, may increase mortality (Lirman, 2000; Forsman et al., 2006) and lead to a loss of fecundity (Harrison & Wallace, 1990; Zakai et al., 2000; Okubo et al., 2007), limiting dispersal and recruitment in reefs restored in this manner until coral colonies reach mature sizes (Kozłowski & Wiegert, 1986; Ward, 1995; Smith & Hughes, 1999). Curiously, micro-fragments of corals display short-term accelerated growth immediately following initial fragmentation and resulting injury to tissues (Forsman et al., 2015; Page et al., 2018). The mechanisms underlying this phenotypic response remain poorly understood. Early studies on corals that focused on recovery from physical injury suggested that energy required to heal lesions was sourced almost exclusively from adjacent cells (Bak & Yvonne, 1980; Bak, 1983; Meesters et al., 1994), which is consistent with a lack of cellular specialization and postulated low levels of colony integration in corals. However, Oren et al. (2012) demonstrated that lesions in the merulinid *Dipsastrea favus* healed more effectively in large colonies compared to smaller ones. Further, even polyps distant from the lesion showed reduced fecundity, indicating that resources for regeneration were translocated across the colony rather than just from cells bordering the lesion. Such findings indicate that coral colonies are more highly integrated than traditionally thought.

Massive *Porites* spp. have a documented high regeneration capacity, which may be related to the integration of resources throughout their often large colonies (e.g., Lough & Barnes, 2000). Indeed, *Porites* spp. have been shown to heal from tissue damage induced by micro-fragmentation within days, initiating calcification along fragment margins within a week (Forsman et al., 2006, 2015). However, the physiological mechanisms that underlie this response, which is tied to the success of restoration efforts, have not been thoroughly explored. Here, we employed a time-series transcriptomic approach to shed light on the cellular-level mechanisms that drive lesion healing and accelerated growth in micro-fragments of *Porites lobata*. Contrary to other stress events that dampen energy production, suppress calcification, and increase mortality in corals (e.g., Desalvo et al., 2008; Császár et al., 2009; DeSalvo et al., 2012; Bay et al., 2013; Tarrant et al., 2014; Maor-Landaw & Levy, 2016; Aguilar et al., 2019), we provide evidence from gene expression profiles that indicate a temporary increase in energy metabolism and a phosphoinositide-mediated disruption of calcium-homeostasis that likely impacts cell cycle regulation and growth. The endoplasmic reticulum (ER) plays an important role in regulating calcium homeostasis and mitigating the impacts that increased mitochondrial reactive oxygen production has on protein synthesis during physiological stress. We provide a model of the cellular processes that are likely driving the increase in growth rates of *Porites* spp. in the days and weeks following micro-fragmentation.

## 2 MATERIALS AND METHODS

### 2.1 Sampling of source colonies

Six *Porites lobata* colonies, ranging in size from 15-25cm, were sourced from the Luminao reef flat, located on the western coast of Guam. All colonies were collected from ∼2 m depth and were at least 20 m apart to minimize the possibility that they were of clonal origin. Colonies were immediately transported in fresh seawater to the Marine Laboratory at the University of Guam and allowed to acclimate for four weeks in a flow-through seawater tank under 70% shade. Oscillating water motion was maintained with OW-40 wavemakers (Zhongshan Jebao Electronic Appliance Co, Beijing, China).

### 2.2 DNA barcoding and algal symbiont profiling

Because *in situ* species-level identification is difficult in massive *Porites* spp., colonies with similar gross morphologies were sampled and species identifications were then verified using DNA barcoding. DNA was extracted using the GenCatch genomic DNA extraction kit (Epoch Life Science, Sugar Land, TX) following the manufacturer’s protocol for tissue samples. Mitochondrial markers mt-16 (Cox3-Cox2) and mt-20 (ND5-tRNA-Trp-ATP8-Cox1) (Paz-García et al., 2016) were amplified via PCR using 0.3uM primer, 0.3mM dNTP, 14.25 uL Water, 14.25ul HiFi Fidelity Buffer (5x), 2.5 units Taq (HIFI kapa 1U) and denaturation at 94°C for 120 s, followed by 30 cycles of 94°C for 30 s, 54°C for 30 s, 72°C for 60 s, with a further extension step of 72°C for 300 s. PCR products were sequenced in both directions using Sanger sequencing and resulting sequences assembled using the overlap-layout-consensus algorithm implemented in Geneious Prime (Biomatters, Auckland, New Zealand). Following assembly, consensus sequences for each specimen were aligned using MUSCLE (v. 3.8; Edgar, 2004) and sequences compared to NCBI’s GenBank nucleotide sequence database.

To determine the dominant clade of Symbiodinaceae associated with each sample, transcriptomic reads from each source colony were mapped against symbiont transcriptomes for the four Symbiodiniaceae genera that associate with scleractinian corals (*Symbiodinium, Breviolum, Cladocopium, Durusdinium*; LaJeunesse et al., 2018). The number of high-quality mapped reads for each Symbiodinaceae genera where divided by the total high-quality mapped reads to determine relative abundance, following the approach of Manzello et al., (2019).

### 2.3 Micro-fragmentation, *ex situ* husbandry, and outplanting

After four weeks of acclimation in flow-through seawater tanks, colonies were cut into ∼1.5 cm^2^ micro-fragments using a seawater-cooled C-40 diamond band saw (Gryphon Corporation, Sylmar, CA), glued onto individual tiles, and randomly assigned to one of two replicate tanks. Micro-fragments were created from coral tissue at least 2 cm from the growing edge of colonies to reduce variability in growth rates and gene expression differences between fragments. Excess skeleton was removed from the base of each fragment to create micro-fragments of the same height, which were then attached to ceramic tiles using cyanoacrylate gel and labeled according to source colonies. In total, 36 fragments were cut (n=6/colony), to be destructively sampled for transcriptomics at predetermined time points and an additional 42 fragments were prepared (n=7/ colony) to document growth rates. After micro-fragments had grown for two months in the two flow-through tanks, 42 micro-fragments were affixed to natural reef substrate using Splash Zone epoxy (Z-spar, Périgny, France), in three plots at Asan Beach National Park, within a Marine Preserve on the western coast of Guam. The outplanting occurred in November, which is outside of the normal bleaching season for Guam.

### 2.4 Growth rates

Weekly maintenance of the micro-fragments in *ex-situ* culture included removal of algae and detritus from tiles and tanks, and visual inspection of all tiles to identify and remove potential nudibranch predators. Water temperature was monitored using HOBO Tidbit temperature loggers (Onset, Bourne, MA, USA). Weekly photographs of each fragment (top-down and side-views from all four sides of each tile) were taken using an Olympus TG-5 (Olympus, Shinjuku, Tokyo, Japan) in an underwater housing mounted to a PVC stand with attached scale bar. Micro-fragment surface area was estimated from photographs using ImageJ version 1.43u (National Institutes of Health, Bethesda, MD), combining surface areas measured from top and side views for each micro-fragment. Growth was determined via the change in surface area from one week to the next and normality of growth rate distributions was tested using a Shapiro-Wilks test.

Growth was then binned into two phases based on a pilot study, Phase A from T1 (week 0) to T3 (week 3) during which cut margins were overgrown by new tissue; and Phase B from T3 (week 3) to T5 (week 8) during which coral tissue and skeleton were deposited onto tiles. To account for the non-independence of time-series, Repeated Measures ANOVA and Student’s t tests were implemented in R studio to test for significance between these two growth phases, as well as differences between colonies and rearing tanks.

### 2.4 Transcriptome sequencing, assembly, and annotation

One micro-fragment per colony was destructively sampled for RNA extraction at five time points (Figure 1): after colony collection and acclimation to experimental tanks but prior to micro-fragmentation (baseline; T1); 24 hr after micro-fragmentation (initial stress; T2); at the first signs of calcification along cut margins (regeneration; T3); at 2 mo of *ex situ* growth (recovery; T4); and one day after outplanting (transplantation stress; T5). Tissue samples from the tank experiment (T1-T4) were placed into sterile flat wire sample bags and immediately flash frozen using liquid nitrogen; tissue samples were stored at -80°C until extraction. Transcriptomic samples from the outplanted fragments (T5) were field-preserved in RNAlater and subsequently stored at -80°C until extraction. Total RNA was extracted using an E.Z.N.A. Plant RNA kit (Omega Bio-Tek, Norcross, GA). DNA was removed from RNA extracts using a DNase 1 digest. After extraction, RNA was quantified fluorometrically using a Qubit RNA HS Assay kit (Life Technologies, Carlsbad, CA) and RNA integrity verified using a BioAnalyzer (Agilent Technologies, Santa Clara, CA). RNA sequencing libraries were constructed using the NEBNext Ultra RNA Library Prep Kit (New England Biolabs, Ipswich, MA) following manufacturer’s protocol. Sequencing libraries were multiplexed and 150bp paired-end reads generated on an NextSeq 500 sequencer (Illumina, San Diego, CA) using the Illumina NextSeq 500/550 High Output Kit v2.5 (Illumina, San Diego, CA).

**Figure 1:**
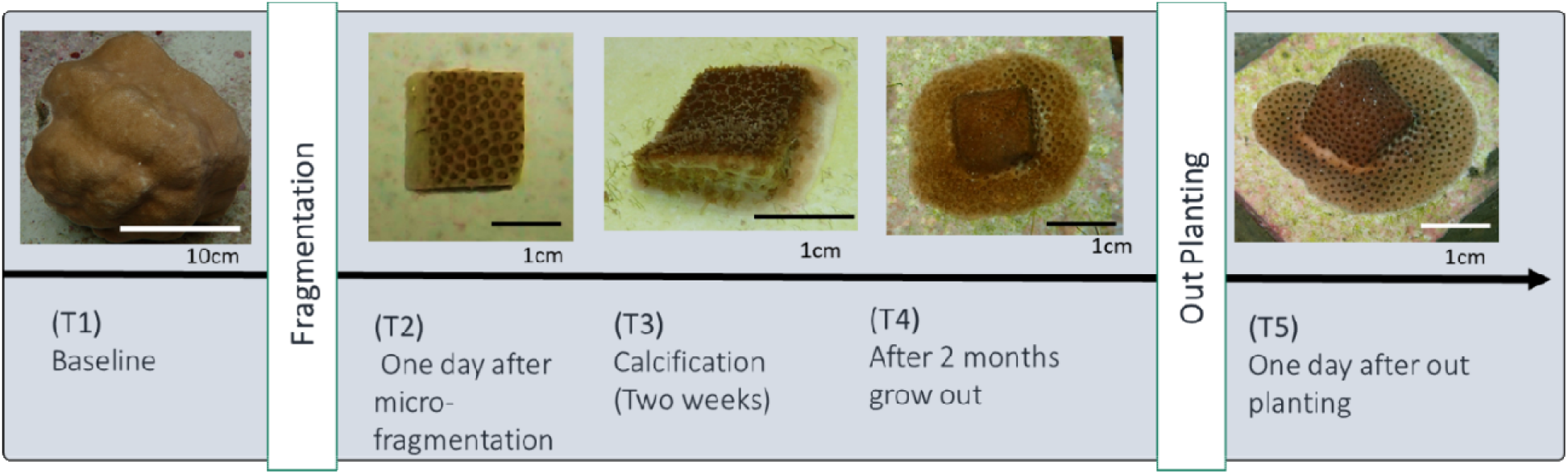
The timeline of transcriptomic sampling.

Adapter sequences and bases with a Phred-scaled quality score of less than 30 were trimmed using Trim Galore (Martin, 2011). Prior to *de novo* transcriptome assembly, all reads were combined and then normalized using in-silico normalization as implemented in Trinity v2.10.0 (Grabherr et al., 2013) with a max coverage set to 50; this normalized read set was then used for transcriptome assembly with Trinity. TransDecoder v5.5.0 (Haas et al., 2013) was used to predict open reading frames (ORFs); transcripts for which no ORF was predicted were removed from further analysis. Proteomes from each major bacterial clade (cf. Schulz et al., 2017), fungi, stramenopiles, poriferans, arthropods, molluscs, and annelids were obtained from NCBI’s GenBank and concatenated into a non-target (alien) BLAST database. Target cnidarian (corals and other cnidarians) and Symbiodinaceae protein sequences were obtained from the Uniprot database. Alien Index (Ryan, 2014) was used in conjunction with protein BLAST searches of our predicted ORFs (proteome) against non-target, alien, and target protein databases to identify and remove likely contaminant sequences from our *de novo* assembled transcriptome. This strategy yielded a meta-transcriptome consisting of coral and Symbiodinaceae transcripts and predicted ORFs. This meta-transcriptome was then annotated with gene ontology (GO) terms (Ashburner et al., 2004) retaining the top hit from BLAST searches (e-value <= 1e-5) against the combined cnidarian and Symbiodinaceae Uniprot database. The reference meta-transcriptome was then parsed into *Porites lobata* host and Symbiodniaceae symbiont transcriptomes using Alien Index with target cnidarian and Symbiodinaceae protein sequences for differential gene analysis.

The Benchmarking Universal SingleCopy Orthologs (BUSCO) (Simão et al., 2015) pipeline was used to determine the completeness of both the *P. lobata* and Symbiodinaceae transcriptomes. BUSCO runs were performed on the transcript level, comparing transcriptome assemblies against the metazoan database for *P. lobata* and alveolates for Symbiodinaceae. In *de novo* transcriptome assemblies, it is common to find several predicted isoforms for the same gene, which can lead to high duplication rates in BUSCO results. To adjust for such artificially inflated duplication rates, transcripts originating from the same gene graph in the assembly that are inferred to originate from the same gene were counted as a single hit in BUSCO.

### 2.5 Differential gene expression and gene ontology

Coral and Symbiont transcript and gene abundances were inferred separately using the fast k-mer hashing and pseudoalignment algorithm implemented in Kallisto (v0.46.2; Pimentel et al., 2017). The R package Sleuth (v. 0.30.0; Pimentel et al., 2017) was used to identify differentially expressed transcripts (*q*-value < 0.01). Significance levels of transcripts belonging to the same gene (see BUSCO analysis under 2.4) were aggregated (*p*-value aggregation method; Pimentel et al., 2017) to infer differential expression at the gene, rather than transcript, level. Time series differential analysis based on natural spline models (i.e., a likelihood ratio test) was used to discern patterns of expression across all time points. A natural spline model (df = 4) was used to fit “knots” along the observations of the time axis (full model) to determine if gene expression followed a non-random pattern compared to a model of random variation (null model). A likelihood ratio test, as implemented in Sleuth, was employed to compare full and null models for each gene to identify differentially expressed genes along the time series.

The Transcripts Per Million (TPM) values of differentially expressed genes were used to produce a heatmap using the pheatmap package (version 1.0.12) in R with scaling set to row (Figure 3). clustered hierarchically using the hclust function in R to identify likely co-expressed genes. The dendrogram of genes (left side of figure 3) was cut at h=9.5 to obtain gene clusters and then visually condensed into patterns that are up or down regulated at similar points in the timeseries. The differentially expressed genes from the major patterns (Figure 4) used for GO enrichment using a Fischer’s exact test in REVIGO version to determine significant up or down regulated GO categories (Supek et al., 2011). In addition, pairwise Wald tests, as implemented in Sleuth, were used to identify differentially expressed genes between time points.

## 3 RESULTS

### 3.1 Coral host and algal symbiont identities

Coral host colonies were found to be the same species based on coral mitochondrial sequences (NCBI Accession nos. OM858841-OM858852), which were invariant across the almost 2,250bp sequenced apart from a single nucleotide polymorphism in specimen five. Sequence similarity searches against NCBI’s GenBank identified the mitochondrial genome of *Porites lobata* (KU572435;Tisthammer et al., 2016) as the most similar publicly available sequence data. Query coverage was 100% for all specimens and loci, with similarities greater or equal to 99.9%. Based on transcript mapping to the putative Symbiodiniaceae transcriptomes, the algal symbionts for all samples were primarily (>93%) from the genus *Cladocopium* (Supplementary Figure 1).

### 3.2 Micro-fragment growth rates

All 42 *P. lobata* micro-fragments survived the *ex-situ* phase of the experiment without signs of disease, predation, or bleaching. On average, micro-fragments increased in surface area by 355.4% over the 8-week duration of the experiment in the flow-through seawater tanks (Figure 2). Growth rates were normally distributed (Shapiro-Wilk’s test; *p*-value = 0.067) and growth rates between replicate tanks were not significantly different (*t=*-0.52893; *p*-value = 0.271). Two distinct growth phases were identified (Repeated Measures ANOVA, df = 92; *p*-value < 0.001): Phase A (weeks 0 – 2) characterized with rapid growth of tissue over cut margins of the micro-fragment and Phase B (weeks 3 – 8) characterized by new growth deposited onto the tile. Significant variation in growth rates during phase B was found between colonies (*p*-value = 0.002), with colonies 1 through 4 continuing to grow rapidly while colonies 5 and 6 showed reduced growth rates after initial wound healing (Figure 2). Two weeks after outplanting, the newly grown tissue of half of all outplanted micro-fragments (16/32) showed signs of bleaching despite growing in a similar temperature regime as that of the *ex-situ* tanks. No signs of disease or predation were observed during the *ex-situ* culture or after outplanting.

**Figure 2:**
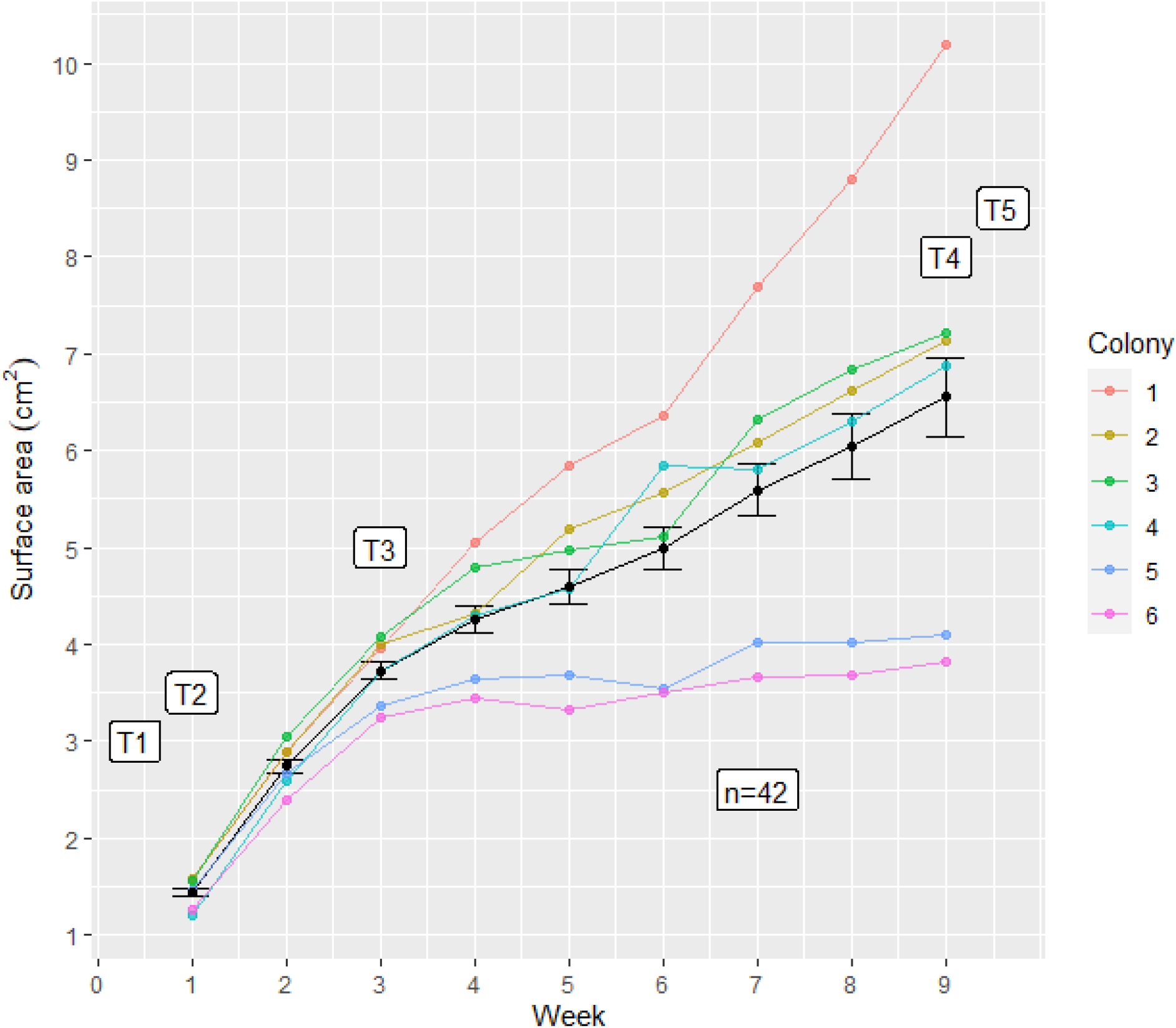
The average (Mean±SE) weekly surface for the 42 *Porites* micro-fragments over the 8-week experiment. T1-T5 represent the transcriptomic sampling points. Colored lines represent individual colony growth, and the black line represents mean micro-frag growth.

### 3.3 Reference assembly

Illumina sequencing generated between 6.9 and 54 million reads per sample with an average of 35 million reads per sample (Supplemental Table 1; BioProject number: PRJNA757218). Due to a low number of reads generated at some timepoints, colony 5 (Supplemental Table 2) was removed from further analysis. Trinity *de novo* transcriptome assembly generated 1,963,624 transcripts, which were filtered and annotated to produce a reference transcriptome composed of 60,475 coral and 17,994 symbiont gene models (Supplementary Table 1). BUSCO analysis indicated that the reference for the coral host was almost complete (C: 93.7%; S: 13.0%; D: 80.7% [adjusted D: 4.5%]; F: 2.5%; M: 3.8%; n: 954) while the symbiont reference was roughly 70% complete (C: 69.6%; S: 39.2%; D: 30.4% [adjusted D:23.4%]; F: 4.1%; M: 26.3%; n: 171) (Supplementary Table 3).

### 3.4 Time series gene co-expression

The separate analysis of symbiont and coral transcripts revealed a total of 2,282 coral genes and 44 symbiont genes were identified as differentially expressed by the spline regression analysis (*p*-value < 0.01; Figure 3). Hierarchical clustering (D’haeseleer, 2005) of significantly differentially expressed genes clustered the samples by time point replicate and then by stress event (T2, T5) vs. non-stress event (T1, T3, T4; columns in Figure 3). The dendrogram of differentially expressed genes (left side of Figure 3; Figure 4a) indicated 26 clusters of genes that were condensed into six major patterns of expression through the time series (Figure 4). Three of these patterns (4-6) consisted of relatively few genes and produced no significant GO enrichment results and were thus removed from further analysis. The remaining three clusters consisted of enriched gene categories from genes that were upregulated at the stress events (T2 & T5; Figure 4b), genes that were downregulated during the stress events (Figure 4c), and genes that were downregulated during the rapid initial Growth Phase A (Figure 4d).

**Figure 3:**
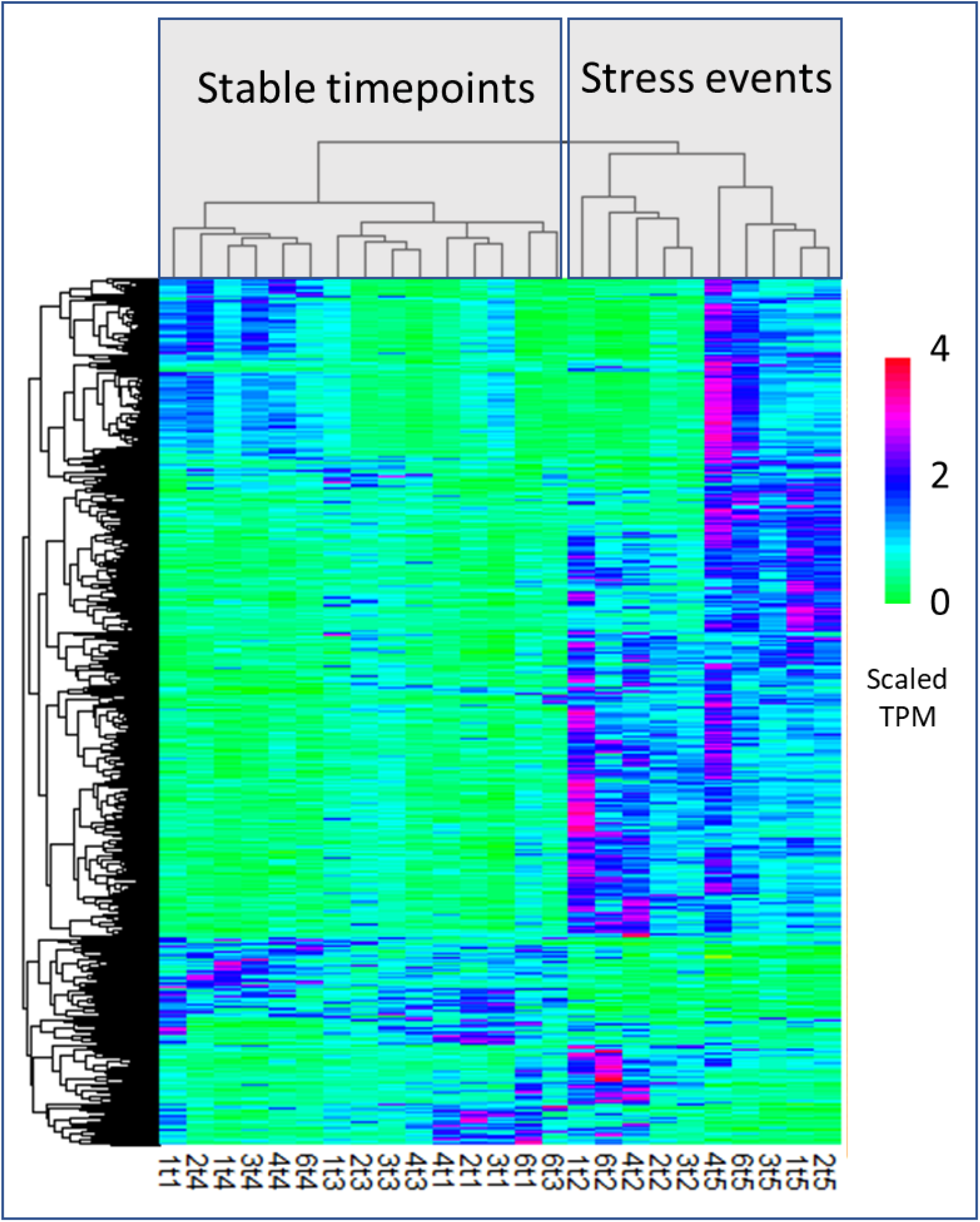
Heat map of significantly differentially expressed coral host genes identified by spline regression analysis. The dendrogram on the left corresponds to clustering of genes (rows) by expression differences across samples (columns). The first number of the column name corresponds to colony replicate and the second refers to the sampling timepoint. The gene expression patterns cluster the samples into two main clusters: the stress events (T2; fragmentation and T5; outplanting) and the more stable growth points (T1, T3, & T4). The color scale represents the Kallisto TPM values scaled by the pheatmap (scale = “row”) package in R.

**Figure 4:**
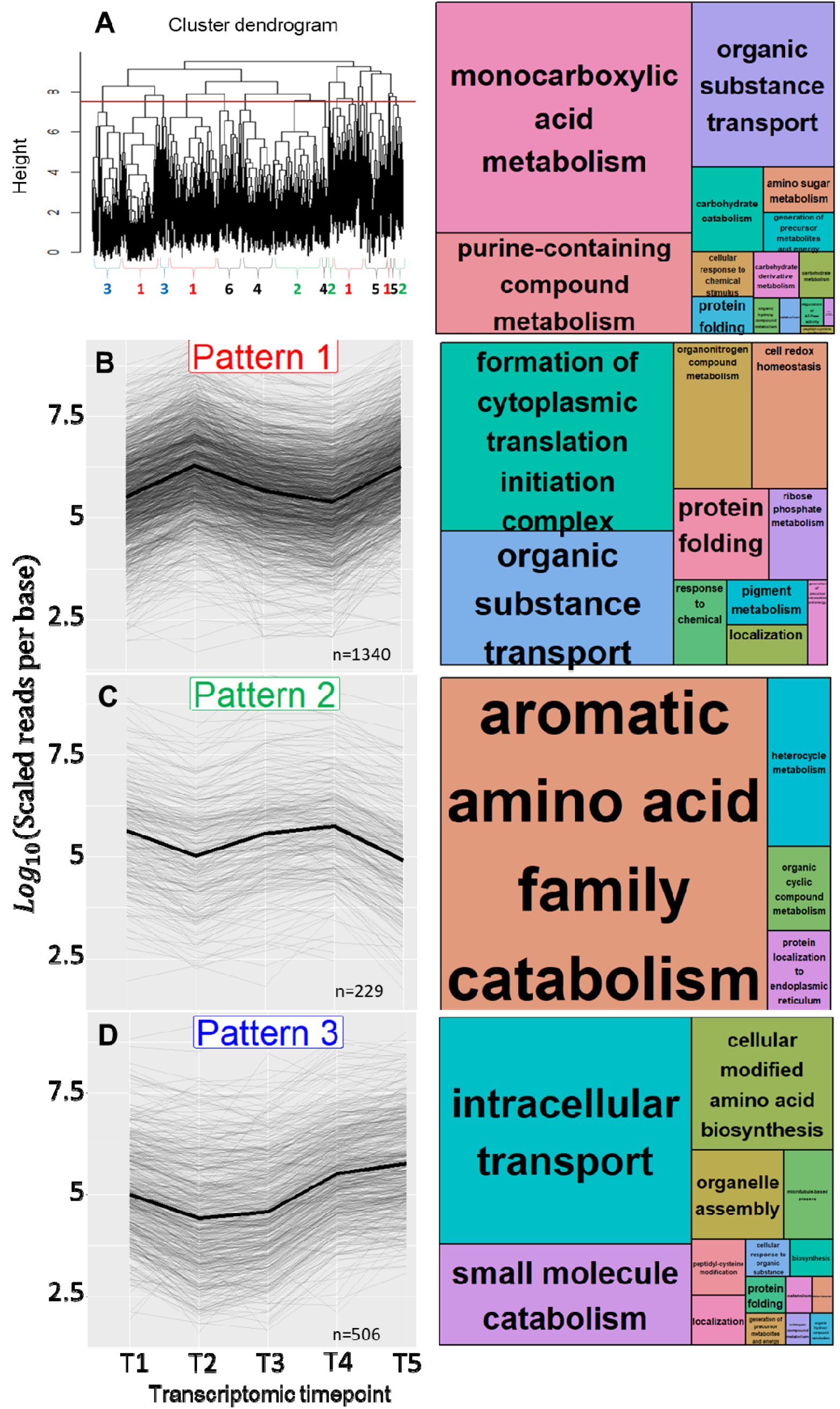
Major patterns of gene expression clusters through the time series with GO enrichment patterns. Upper left (4a) is the cluster dendrogram from the heatmap (Figure 3) with the red line defining the cutting of gene clusters. The upper right figure shows REVIGO GO enrichment treemap of all significant genes (n=2286). The three major gene patterns (4b; pattern 1 = 1340 genes, 4c; pattern 2 = 229 genes, 4d; pattern 3 = 506 genes) identified through the time series with their corresponding REVIGO GO enrichment treemaps are presented here. Text size correlates with the significance (large text = smaller pval) of enriched gene category. See supplemental text for a complete list of enriched GO categories.

GO enrichment of all identified significant genes (n = 2286) across the time series yielded terms associated with carbohydrate metabolic process (GO:0005975), phosphorus metabolic process (GO:0006793), small molecule metabolic process (GO:0044281), proteolysis (GO:0006508), oxidation-reduction process (GO:0055114), nucleobase-containing compound biosynthetic process (GO:0034654), organophosphate metabolic process (GO:0019637), phosphate-containing compound metabolic process (GO:0006796), phosphorylation (GO:0016310), organic cyclic compound biosynthetic process (GO:1901362), aromatic compound biosynthetic process (GO:0019438), heterocycle biosynthetic process (GO:0018130), ion transport (GO:0006811), organonitrogen compound biosynthetic process (GO:1901566), cellular nitrogen compound biosynthetic process (GO:0044271), transmembrane transport (GO:0055085), organic acid metabolic process (GO:0006082), small molecule biosynthetic process (GO:0044283), carboxylic acid metabolic process (GO:0019752), cellular amino acid metabolic process (GO:0006520), and oxoacid metabolic process (GO:0043436).

GO categories that were significantly upregulated at both of the stress events (Pattern 1, 1340 genes) include formation of cytoplasmic translation initiation complex (GO:0001732), protein folding (GO:0006457), cell redox homeostasis (GO:0045454), generation of precursor metabolites and energy (GO:0006091), ribose phosphate metabolism (GO:0019693), pigment metabolism (GO:0042440), organonitrogen compound metabolism (GO:1901564), response to chemical (GO:0042221), localization (GO:0051179), and organic substance transport (GO:0071702). GO categories that were significantly downregulated at both stress events (Pattern 2, 229 genes) include aromatic amino acid family catabolism (GO:0009074), heterocycle metabolism (GO:0046483), organic cyclic compound metabolism (GO:1901360), and protein localization to endoplasmic reticulum (GO:0070972).

GO categories that were significantly downregulated when the micro-frags were recently cut and rapidly growing (Pattern 3, 506 genes) include intracellular transport (GO:0046907), cellular modified amino acid biosynthesis (GO:0042398), small molecule catabolism (GO:0044282), microtubule-based process (GO:0007018), localization (GO:0051179), peptidyl-cysteine modification (GO:0018198), cellular modified amino acid biosynthesis (GO:0042398), protein folding (GO:0006457), catabolism (GO:0009056), generation of precursor metabolites and energy (GO:0006091), organelle assembly (GO:0070925), cellular response to organic substance (GO:0071310), nitrogen compound metabolism (GO:0006807), and organic hydroxy compound metabolism (GO:1901615).

### 3.5 Pairwise gene expression analysis

While analyzing coral host gene expression profiles during the transition between states, the largest number of differentially expressed genes was identified immediately following fragmentation (T1 versus T2) (Table 1). The transition between T2 and T3 yielded few differentially expressed genes. Additionally, the comparison between the baseline (T1) and two weeks of growth (T3) yielded few differentially expressed genes. The response to fragmentation (T1 versus T2) yielded twice as many differentially expressed genes as the response to outplanting (T4 versus T5). Previous literature on the Cnidarian stress response was used to identify crucial metabolic genes/pathways and those genes found differentially expressed in this study are listed in Supplemental Table 4. Several crucial antioxidant enzymes were upregulated in response to stress in this study, including Thioredoxin, Peroxiredoxin-1, Peptide-methionine (S)-S-oxide reductase (EC 1.8.4.11), and Glutathione transferase (EC 2.5.1.18) (Supplemental Table 4). Heat shock proteins were upregulated in response to fragmentation and outplanting, such as HSP70-1, HSP90, 97 kDa heat shock protein, HSP40s, and 71kDa heat shock proteins (Supplemental Table 4). A variety of protein degradation and biosynthesis enzymes were identified as differentially expressed in response to both stress events (Supplemental Table 4), including Ubiquitin-associated ligases, hydrolases, conjugation factors, and transferases, translation initiation factors, ribosomal biogenesis proteins, tRNA ligases, deacylases, synthetases, and hydrolases. Calcium homeostasis genes were significantly upregulated in response to fragmentation, such as Calreticulin, calumenin, Voltage-dependent L-type calcium channel subunit alpha, Calcium-transporting ATPase, and many genes which cause the release of calcium into the cytosol via the phosphatidylinositol signaling pathway. Energy metabolism genes relating to glycolysis, gluconeogenesis, the citric acid cycle, and electron transport chain were upregulated immediately following fragmentation (Supplemental Table 4).

**Table 1:** The number of differentially expressed genes between all pairwise timepoints for coral host and symbiont.

| **Coral genes** | T1(Baseline) | T2(Immediate response) | T3 (2 full weeks of growth) | T4 (2 months of growth) |
| --- | --- | --- | --- | --- |
| T1(Baseline) |  |  |  |  |
| T2 (Immediate response) | 1648 |  |  |  |
| T3 (2 full weeks of growth) | 73 | 64 |  |  |
| T4 (2 months of growth) | 116 | 1794 | 2 |  |
| T5 (Outplanting) | 1753 | 1207 | 299 | 820 |
| ***Symbiodinium* genes** |  |  |  |  |
| T1 (Baseline) |  |  |  |  |
| T2 (Immediate response) | 7 |  |  |  |
| T3 (2 weeks of growth) | 0 | 7 |  |  |
| T4 (2 months of growth) | 2 | 42 | 5 |  |
| T5 (Outplanting) | 11 | 210 | 33 | 2 |

#### Oxidative stress

Antioxidant molecules reduce reactive oxygen species, oxidize proteins, and stabilize proteins through cellular stress (Heat shock proteins, components of the thioredoxin system). Several crucial antioxidant enzymes were upregulated in response to stress in this study (Supplemental Table 4). Thioredoxin, Peroxiredoxin-1, Peptide-methionine (S)-S-oxide reductase (EC 1.8.4.11), and Glutathione transferase (EC 2.5.1.18) were upregulated in response to fragmentation and outplanting Glutathione peroxidase was upregulated in response to outplanting only. No antioxidant-related genes were found upregulated after two weeks (T3) or two months (T4) of micro-fragment growth. Ferritin (EC 1.16.3.1), an iron-binding protein that controls the amount of available ferrous iron (Fe^2+^), which is involved in free radical generation, was upregulated after fragmentation and outplanting. Hypoxia up-regulated protein 1 was upregulated after fragmentation and outplanting. Thioredoxin, Glutathione transferase, and Glutathione peroxidase were not identified as differentially expressed after two weeks (T3) and two months (T4) of growth.

Heat shock proteins (HSPs), which serve a variety of protein stabilizing and folding functions in response to stress, were found upregulated in response to immediate fragmentation and outplanting (Supplemental Table 4). HSP70-1 was upregulated after the immediate fragmentation and remained upregulated after two weeks of rapid growth. Four co-chaperones (DnaJ-like proteins) were upregulated in response to fragmentation. Two DnaJ-like proteins were upregulated in response to outplanting. Interestingly, after two weeks (T3) and two months of growth (T4), no HSPs were differentially expressed, indicating a potential return to protein stability. Five out of six HSPs identified be significantly upregulated in response to fragmentation. Six out of seven HSPs be upregulated after the fragments were outplanted.

#### Protein degradation, synthesis, and transport

Stress events cause a breakdown of normal protein homeostasis leading to increased degradation, synthesis, and transport of proteins (Maor-Landaw & Levy, 2016). A variety of protein degradation enzymes were identified in response to each stress event (Supplemental Table 4). Ubiquitin-associated ligases, hydrolases, conjugation factors, and transferases were differential expressed in response to both fragmentation and outplanting. Eight out of twelve and sixteen out of twenty-two ubiquitin-associated proteins were upregulated after fragmentation (T2) and outplanting (T5), respectively, indicating increased rates of protein degradation in response to these stress events. Only one Ubiquitin-associated protein was downregulated at either fourteen days (T3) or two months after fragmentation (T4), which likely demonstrates that the increase in protein degradation processes returns to baseline expression levels quickly following these stress events. Three out of four protein disulfide-isomerases and two 26s proteasome regulatory subunits were upregulated in response to fragmentation (T2).

Several genes associated with protein anabolism were found differentially expressed in response to the stress events, indicating significant increase in protein synthesis. A total of 17 out of 17 translation initiation factors were upregulated in response to fragmentation. Three ribosomal proteins (Ribosome production factor 2 homolog (Ribosome biogenesis protein RPF2 homolog), Ribosomal protein S6 kinase (EC 2.7.11.1), and Ribosomal RNA small subunit methyltransferase NEP) associated with protein biogenesis were upregulated in response to outplanting (T5). In addition, a variety of tRNA ligases, deacylases, synthetases, and hydrolases were all upregulated in response to fragmentation (15/15) and in response to outplanting (5/5). Four out of five aminotransferase enzymes were found upregulated in response to outplanting (T5). 5-aminolevulinate synthase was upregulated in response to fragmentation (T2). Amino acid transporters were found upregulated in response to fragmentation (3/3) and outplanting (1/1).

#### Cell Cycle

Interestingly, four growth factors and receptors were downregulated in response to fragmentation (T2), when the corals began their most rapid growth phase (Supplemental Table 4; Figure 5). Fibroblast growth factor receptor 1-A was significantly upregulated after two weeks of growth (T3), relative to in response to fragmentation. Important apoptosis-inducing genes (caspase-3, programmed cell death protein 6, and apoptosis regulator BAX) were upregulated in response to both fragmentation (T2) and outplanting (T5). In contrast, Caspase-7 was upregulated only in response to outplanting, whereas a bifunctional apoptosis regulator was upregulated after only fragmentation.

**Figure 5:**
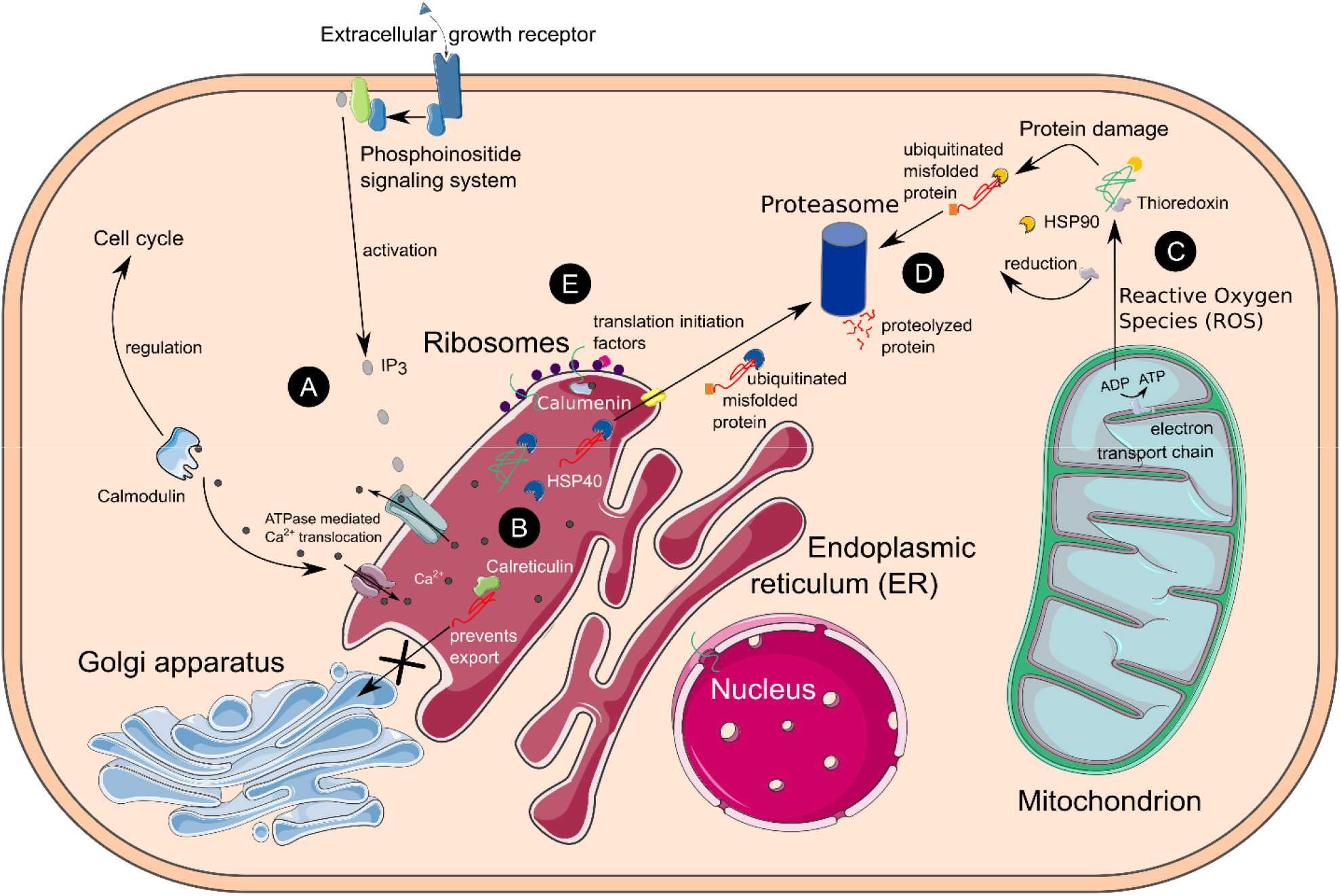
Cell diagram of the integration of calcium homeostasis disruption (A), Endoplasmic-Reticulum (ER) stress (B), protein anabolism (D) increased energy production (C), and protein degradation (D) found upregulated 24 hours after physical injury. 5A: Phosphoinositide signaling releases the secondary intracellular messenger, inositol (1,4,5) trisphosphate (IP3), which binds to ligand-gated calcium ion channels of the ER causing the release of calcium into the cytosol. Calcium disruption is sensed by a variety of molecules including calmodulin (upregulated in this study), which leads to enzymatic activation and ultimately, cell cycle regulation. Calmodulin also stimulates the reuptake of Ca^2+^ into the ER to prevent prolonged calcium homeostasis disruption and cell death. **5B:** Calcium disruption within the ER leads to the Unfolded Protein Response (UPR). Resident proteins upregulated in this study, such as calreticulin, calumenin, and Heat Shock Proteins (HSPs), assist in the folding of proteins and degradation of terminally misfolded proteins. Calreticulin prevents the export of misfolded proteins to the golgi apparatus. **5C:** Ribosomal proteins, translation initiation factors, and tRNA enzymes were significantly upregulated and rapidly produced new proteins in response to fragmentation. **5D:** To meet energetic demands of the cell under stress, expression of electron transport chain proteins increased. Increased oxidative phosphorylation causes Reactive Oxygen Species (ROS) to leak from the mitochondria, causing cellular damage to lipids, DNA, and proteins. Antioxidant molecules upregulated in this study (thioredoxin, glutathione transferase, peroxiredoxin, ferritin) scavenge ROS and assist in protein refolding. Cytosolic HSPs refold damaged proteins and assist in their degradation if they are terminally misfolded. **5E:** Terminally misfolded proteins are chaperoned to the proteasome by HSPs after they are tagged for destruction by Ubiquitin-conjugating enzymes. The Ubiquitin/Proteosome system integrates with other signal transduction molecules to regulate cell cycle. Amino acid transporters shuttle degraded polypeptides to ribosomes for protein anabolism.

#### Cytoskeleton and extracellular matrix

Cytoskeleton rearrangement is often required to maintain cell motility, structure, and integrity in response to cellular stress. Several cytoskeleton and extracellular matrix genes were differentially expressed in response to the stress events (Supplemental Table 4). Carbonic anhydrase (EC 4.2.1.1), which is understood to be important in coral carbonate deposition (Maor-Landaw & Levy, 2016), and beta-actin was found downregulated whereas three alpha tubulin proteins were upregulated in response to fragmentation (T2). Alpha tubulin was also upregulated in response to outplanting (T5).

#### Calcium

Calcium acts as an intracellular secondary messenger and mediates a variety of function within the cell. Several genes related to calcium homeostasis disruption were differentially expressed (Supplemental Table 4). Calrecticulin, calumenin, Voltage-dependent L-type calcium channel subunit alpha, and Calcium-transporting ATPase (EC 7.2.2.10) were significantly upregulated in response to fragmentation (T2). Calmodulin, which plays an important role in calcification and calcium signaling, was upregulated in response to fragmentation, downregulated after 2 weeks of growth (T3), and not differentially expressed after two months of growth (T4). Calcium and integrin-binding family member 2 was upregulated in response to outplanting. Guanylate cyclase, which is involved in energy conversion, G protein signaling cascade, and is inhibited by high intracellular calcium levels, was downregulated in response to outplanting.

#### Lipid metabolism

Eleven genes relating to the phosphatidylinositol signaling pathway, which stimulates the release of calcium from the endoplasmic reticulum, were upregulated in response to fragmentation (T2), whereas four out of six were upregulated in response to outplanting (T5). No phosphatidylinositol signaling-related genes were differentially expressed after two weeks or after two months of growth. Sphinogolipid 4-desaturase was found upregulated in response to fragmentation and in response to outplanting. Two acyl-coenzyme A thioesterases and Mitochondrial carnitine/acylcarnitine carrier protein CACL, which is involved in lipid catabolism, transport, and signaling, were upregulated, and an O-acyltransferase was found downregulated, in response to fragmentation. Lipid droplet-associated hydrolase, Very-long-chain 3-oxoacyl-CoA synthase (EC 2.3.1.199), and 3-hydroxyacyl-CoA dehydrogenase type-2 were upregulated in response to outplanting.

#### Carbohydrate metabolism

A variety of carbohydrate metabolism genes were found upregulated in response to the two stress events. Phosphoglycerate mutase (EC 5.4.2.12), which is involved in glycolysis, was upregulated in response to fragmentation (T2). Phosphoenolpyruvate carboxykinase (EC 4.1.1.32), an enzyme essential for gluconeogenesis, was upregulated in response to fragmentation. Several enzymes involved in the citric acid cycle were upregulated after fragmentation, including ATP-citrate synthase (EC 2.3.3.8), Aconitate hydratase, Mitochondrial pyruvate carrier, and Succinate--CoA ligase (EC 6.2.1.4), and Malate dehydrogenase (EC 1.1.1.37), Mitochondrial pyruvate carrier, and Succinate dehydrogenase (EC 1.3.5.1), in response to outplanting.

#### Cellular energy

Responding to environmental stress is an energetically costly processes that must be met with increased energy production for an organism to persist. A total of 36 crucial metabolism genes involved in the electron transport chain and oxidative phosphorylation were differentially expressed in response to fragmentation (T2) or outplanting (T5), indicating significant energy generation in response to the stress events. (Supplemental Table 4). ATP synthase subunits alpha, beta, B1, and gamma were upregulated in response to fragmentation. Adenylate kinase (EC 2.7.4.3), essential to cellular energy homeostasis by converting between various adenosine phosphates (ATP, ADP, AMP), was upregulated in response to fragmentation. Ten subcomplexes of NADH dehydrogenase, five subcomplexes of cytochrome c oxidase, two subcomplexes of Cytochrome b-245, three subcomplexes of cytochrome b-c1, cytochrome b5 and NADPH cytochrome p450 reductase were all upregulated in response to fragmentation, indicating significant energy generation. Two subunits of cytochrome p450 were the only cytochromes that were downregulated in response to the two stress events. Cytochrome b-561 and ATP synthase subunit alpha were the only parts of the electron transport chain upregulated in response to outplanting.

## 4 DISCUSSION

Growth and calcification rates are negatively correlated with size in corals (Chadwick-Furman et al., 2000) and both increase in response to fragmentation (Lirman et al., 2010; Forsman et al., 2015). Rapid regeneration is crucial to coral colony survival, as it prevents possible infections by pathogens and damage from competitors, and ensures the physical integrity of the coral colony by cementing fragments to each other and the substrate after breakage or fracture. Here, we show that fragmentation causes significant differential expression of genes associated with a disruption in calcium homeostasis and initiates rapid wound healing and recovery in *Porites lobata*. The fragmentation response is characterized by two distinct phases: (1) rapid tissue regeneration at wound margins, followed by (2) a slower growth phase during which the colony is stabilized by new tissue deposited onto the substrate (Figure 2). The initial increase in growth rate following fragmentation of coral colonies has been demonstrated previously (Page et al., 2018), but the underlying cellular-level processes driving the accelerated growth have remained largely unexplored and are crucial for understanding coral regeneration.

The initial stress response to both outplanting and fragmentation yielded specific genes (Supplemental Table 4) and GO terms (Figure 4a) associated with ER stress, protein turnover, and antioxidant production, which is consistent with a generalized stress response seen in other cnidarians (Maor-Landaw & Levy, 2016). However, fragmentation initiated some unique processes, such as upregulation of phosphoinositide-mediated acute calcium homeostasis disruption and a variety of electron transport genes, which we believe stimulate (Fenteany et al., 2000; Mosblech et al., 2008) and provide immediate energy for the rapid growth of the coral colony (Osinga et al., 2011) in a push for wound healing, and ultimately, survival.

Overall, the immediate fragmentation response was characterized by the largest number of differentially regulated genes (1648; *q*-value = 0.01) in our experiment, indicating that fragmentation elicits a series of immediate and unique physiological responses (Table 1). During this initial phase, the margins of the fragment were sealed quickly by new tissue with growth slowing once margins were sealed. Concomitantly with this slowing in growth, gene expression stabilized with only two genes differentially expressed when comparing the two-week time point (T3) to the two-month time point (T4), as well as the baseline (T1) to T3 (Table1). We attribute this finding to the recovery of the coral host from the initial stress response to fragmentation and an indication of the return to metabolic homeostasis during this period.

### 4.1 Stress response and energy metabolism

When organisms are presented with environmental stress, the balance between ROS production and antioxidant defenses is disrupted. This frequently leads to an increase in production of antioxidant compounds to prevent significant cellular damage to lipids, proteins, DNA and, ultimately, cell death via apoptosis (Gorman et al., 1999; Scandalios, 2002). Energy generation is crucial in providing necessary resources to enact a shift in metabolism and survival in stressful conditions. However, many cnidarian transcriptomic studies find decreases in energy generation during stress (Veal et al., 2002; Császár et al., 2009; Tarrant et al., 2014; Maor-Landaw & Levy, 2016; Aguilar et al., 2019), which may be related to the breakdown of symbiosis. In contrast, both our spline regression analysis (Figures 3 and 4) and pairwise comparisons of the coral host (Figure 5 & Supplemental Table 4) provide significant evidence of increased cellular energy generation (genes associated with oxidative phosphorylation, electron transport chain cytochromes, glycolysis, and citric acid cycle enzymes; Supplemental Table 4) and lack a detectable gene expression response in the symbiont following coral fragmentation. The maintenance of symbiosis and continued energy production likely provide the resources necessary for antioxidant defenses, cellular protein maintenance, and rapid wound healing (Figure 5). Many of the metabolic pathways identified as differentially expressed in this study (misfolded protein response, HSPs, cell-redox homeostasis, cytoskeleton rearrangement, and calcium homeostasis disruption) coincide with the generalized oxidative stress response described for cnidarians regardless of the type of stress (Veal et al., 2002; Császár et al., 2009; Tarrant et al., 2014; Maor-Landaw & Levy, 2016; Aguilar et al., 2019). However, fragmentation of *P. lobata* appears to elicit several unique responses, such as phosphoinositide-mediated acute calcium homeostasis disruption, and energy production.

Our study found several antioxidant molecules (heat-shock proteins, thioredoxins, ferritin) (Supplemental Table 4), as well as GO categories associated with cell-redox homeostasis (Figure 4b; Pattern 1), were upregulated in response to both fragmentation and outplanting. The upregulation of genes that participate in the thioredoxin oxidoreductase system (thioredoxin, peroxiredoxin-1, Peptide-methionine (S)-S-oxide reductase (EC 1.8.4.11), and glutathione transferase), ferritin, and ion transporters in response to both fragmentation and outplanting further suggest significant increase in antioxidant and protein repair mechanisms to combat oxidative damage from increased energy production necessary for wound regeneration and survival.

Antioxidant molecules that were once thought of as unique biomarkers for heat stress, such as heat-shock proteins (HSPs), are now understood to be upregulated in response to a variety of stressors, including physical injury in this study. Several studies have suggested that the absence of a HSP response is linked with the initiation of apoptosis pathways (Feder & Hofmann, 1999; Gorman et al., 1999; Samali et al., 1999). Thus, the presence of upregulated HSPs indicates a push for survival. In response to physical injury, *P. lobata* colonies demonstrated a diverse upregulation of mitochondrial, cytosol, and ER resident HSPs (HSP70, HSP90, 95kDA HSP, 10kDA HSP, DnaJ-like HSP40s). Arguably the most studied and ubiquitous HSP, HSP70 (Samali et al., 1999), was the strongest upregulated oxidative stress-associated gene in response to fragmentation (log_2_FC 2.95; FDR 1.2e^--9^). The upregulation of 97 kDA HSP in response to fragmentation and subsequent downregulation after two weeks of growth may suggest that oxidative damage to proteins begins to subside within two weeks following physical injury. Additionally, no antioxidant enzymes were upregulated after either two weeks or two months of growth, suggesting that the increased ROS, and therefore antioxidant demands, diminished within a few weeks of physical injury. Although the thioredoxin system and HSPs remain the most studied and indicative biomarkers for cellular oxidative stress, further study is required to understand their specific roles in protein folding, assembly, regulation, and degradation in cnidarians (Feder & Hofmann, 1999; Maor-Landaw & Levy, 2016).

### 4.2 Endoplasmic reticulum stress response and protein turnover

The endoplasmic reticulum (ER) plays crucial roles in controlling protein quality, facilitating the degradation of misfolded proteins and sensing homeostasis changes such as the release of Ca^2+^ into the cytosol. When significant protein damage and calcium homeostasis disruption occur, ER stress pathways are stimulated leading to increased protein stabilization, degradation, and synthesis required to preserve cellular function (Bahar et al., 2016). Significant ER stress during both fragmentation and outplanting was indicated by the upregulation of GO terms associated with protein turnover (protein catabolism, anabolism, folding) (Figure 4b), chaperones involved in protein stabilization and refolding (HSPs, thioredoxin, calreticulin and calumenin), and ubiquitin-associated degradation enzymes (Supplemental Table 4). We found that GO terms associated with amino acid catabolism (Figure 4b) were downregulated in response to fragmentation and outplanting, which suggests that amino acids were preserved to provide the necessary building blocks for increased protein anabolism as corals respond to stress events (Klasing, 2009).

Interestingly, we did not find any evidence for the upregulation of specific ER-stress signal transducers that have been described from other organisms (i.e., inositol-requiring protein-1 (IRE1), activating transcription factor-6 (ATF6), or protein kinase RNA (PKR)-like ER kinase (PERK); Ron & Walter, 2007). Similar to calcium homeostasis disruption (see below), the duration and severity of ER stress determines if an adaptive survival response persists or apoptosis is initiated (Szegezdi et al., 2006; Bahar et al., 2016). The fact that ER-stress signal transducers were not upregulated one day after outplanting suggests that the fragments were already recovering from initial transplantation shock and stress. Decreased rates of translation initiation is one of the earliest stages of significant ER stress, yet we found significant upregulation of translation initiation factors (17/17 identified) one day after fragmentation (Supplemental Table 4). This suggests that corals were recovering from initial ER stress as early as one day after physical injury, upregulating antioxidant defenses, protein stabilizing molecules, and, ultimately, rapid wound regeneration in a push for survival.

### 4.3 Calcium homeostasis disruption and its role in rapid regeneration

Calcium signaling pathways are ubiquitous signal transduction systems, which regulate a broad variety of cellular processes, such as metabolism, apoptosis, cell proliferation, cell to cell communication, gene expression, and secretion (Stefan, 2020). Many cnidarian stress experiments report calcium homeostasis disruption, but few have elucidated the cell signaling mechanisms causing this disruption (Pinzón et al., 2015; Oakley et al., 2017; Aguilar et al., 2019). Among signaling molecules, membrane-bound phospholipids of the phosphoinositide family play key roles in regulating the activity of proteins within the cell (reviewed in Falkenburger et al., 2010). Phosphoinositide signaling networks are composed of a series of transmembrane G-coupled proteins that regulate diverse functions in the cell, including the release of Ca^2+^ from the ER via the binding of inositol 1,4,5-trisphosphate (IP_3_) to ligand-gated ion channel receptors in the membrane of the ER (Berridge & Irvine, 1989; Stefan, 2020).

Although well characterized in other organisms, the specific roles of phosphoinositide signaling in cnidarians remain uncharacterized and the differential expression of these complexes have been reported rarely (Oakley et al., 2017; Aguilar et al., 2019). For example, phosphoinositide phospholipase C, a crucial enzyme that catalyzes the release of IP_3_, was downregulated in response to thermal stress in the anemone model system *Aiptasia* (Oakley et al., 2017). We found that fragmentation caused the upregulation of rate-limiting components of the phosphoinositide signaling pathway and several growth receptors known to stimulate this pathway, including Phosphoinositide phospholipase C and tyrosine kinase growth factor receptors (Supplemental Table 4). We hypothesize that this response was related to the onset of rapid growth and increased calcification rates following fragmentation.

While initial calcium homeostasis disruption can stimulate survival pathways, sustained levels of calcium in the cytosol lead to the initiation of apoptosis (Orrenius et al., 2003; Bagur & Hajnóczky, 2017). Voltage-dependent and ATPase calcium pumps maintain cytosolic Ca^2+^ levels by sequestering calcium in the ER or secreting it extracellularly. We found the genes coding for calcium ATPase pumps upregulated in response to fragmentation, concomitant with calcium homeostasis disruption genes belonging to the phosphoinositide signaling pathway (Supplemental Table 4). The concurrent upregulation of calcium-releasing mechanisms (phosphoinositide signaling) and calcium-dependent ATPase pumps immediately following fragmentation during a period of accelerated growth (Figure 5) suggests that calcium homeostasis is acutely disrupted by physical injury, stimulating cell proliferation and wound regeneration. To promote cell proliferation rather than trigger apoptosis, calcium is released into the cytosol, likely via the phosphoinositide signaling pathway, to stimulate calmodulin-mediated control of the cell cycle. Excess cytosolic Ca^2+^ is then translocated against a gradient using ATPase pumps to restore calcium levels in the ER lumen (Figure 5).

Ca^2+^ increases in the cytosol are sensed by various Ca^2+^-binding proteins and we found that the following genes encoding for Ca^2+^-binding proteins were differentially expressed: calmodulin (CaM), calreticulin, and calumenin (Supplemental Table 4). Ca^2+^-binding proteins are well-known for their role in initiating signaling cascades that ultimately lead to shifts in metabolism. Contrary to the findings of other cnidarian stress experiments (e.g., DeSalvo et al., 2010; Aguilar et al., 2019), CaM was upregulated in response to fragmentation and subsequently downregulated after two weeks of rapid growth. This corroborates our hypothesis that Ca^2+^ concentrations were returning to stable levels in a push for cellular survival. We suggest that CaM was responsible for the initial calcium homeostasis disruption that triggered rapid regeneration after injury by fragmentation in our experiment. Interestingly, CaM has been identified as a regulator of coral larval settlement and metamorphosis (Reyes-Bermudez et al., 2012, 2016), a period in early development characterized by rapid growth.

In addition, we found that the genes coding for the ER-resident proteins calreticulin and calumenin which, together with HSP40 (DnaJ chaperone), prevent the export of misfolded proteins, were upregulated in response to fragmentation (Supplemental Table 4). Increased mitochondrial ROS production (see *stress response and energy metabolism* above) increases oxidative stress that may disrupt protein folding processes, elevating ER stress. We identified the mechanisms that mitigate the impacts of elevated ROS production caused by increased metabolic activity in corals following fragmentation. In particular, calreticulin prevents the export of misfolded proteins from the ER to the Golgi apparatus. Instead, misfolded or damaged proteins are exported to proteasomes for degradation and recycling. Taken together, our gene expression results put the ER at the center of CaM-mediated cell cycle modifications and mitigation of ROS stress impacts on protein synthesis in the aftermath of fragmentation (Figure 5).

### 4.4 Impacts of physical injury on the host-symbiont relationship

Fewer than ten genes were differentially regulated in the zooxanthellate symbiont community between time points (Table 1). The absence of differentially expressed symbiont antioxidant genes suggests that ROS are unlikely to affect the endosymbionts during coral regeneration. Additionally, we found no evidence of nitric oxide (NO) homeostasis breakdown within the host’s response to physical fragmentation, which has been previously correlated with the breakdown of symbiosis in response to coral bleaching (Perez & Weis, 2006; DeSalvo et al., 2010; Maor-Landaw & Levy, 2016). Together, these results suggest that the coral-zooxanthellae symbiosis is far less affected by physical injury than it is by other environmental stressors. Continued efficacy of the host-symbiotic relationship may help meet the energetic needs of the coral host for cellular protein maintenance, antioxidant defense mechanisms, and rapid wound regeneration in response to physical injury.

### 4.5 Physiology of fragmentation response

This study provides a model of proposed physiological mechanisms that enable rapid recovery and stabilization of corals in response to physical injury (Figure 5). The response of *Porites lobata* to fragmentation and outplanting displays many similarities with the response of corals to other stress events, but we documented several important differences. Environmental stress events usually lead to decreased energy production (Maor-Landaw & Levy, 2016), reduced calcification rates (Anthony et al., 2008; Crook et al., 2013; D’Olivo & McCulloch, 2017), and increased mortality (Glynn, 1990) in corals. By contrast, we found that the acute response to physical injury by fragmentation was characterized by upregulation of genes linked to increased energy production. Further, phosphoinositide-mediated calcium homeostasis disruption and ER stress led to the increased expression of antioxidant defense mechanisms and increased rates of protein turnover. Phosphoinositide-mediated acute calcium homeostasis disruption is a likely key mechanism stimulating the wound healing and recovery processes that are evident from the rapid growth of fragmented corals documented here and by others previously (e.g., Page et al., 2018). The severity and length of ER stress and calcium homeostasis disruption often determines the fate of cells in response to environmental change: survival or initiation of apoptosis (Orrenius et al., 2003; Bahar et al., 2016). The concurrent upregulation of calcium disruption and sequestering signals, translation initiation factors and ER stress-associated molecules (protein chaperones and ubiquitin-degradation enzymes), as well as heat shock proteins, one day after physical injury indicates that fragmented corals push for survival and recovery very quickly following injury. In our experiment, this rapid immediate response ceased after two weeks when the exposed margins of the coral skeleton were sealed again with new tissue. Given these results, our recommendation is to provide micro-fragments a recovery period of at least several weeks until cut margins are overgrown with new tissue. All of these responses are consistent with a life history strategy that utilizes fragmentation as a means of asexual propagation and broaden our understanding of this fundamental cnidarian process.

## ACKNOWLEDGEMENTS

This work received support from the University of Guam Sea Grant Master’s Thesis Research Supplemental Grants and the National Science Foundation’s Established Program to Stimulate Competitive Research (OIA-1457769 and OIA-1946352).

## AUTHOR CONTRIBUTIONS

CL, BB, & LJR conceived the project. CL performed the experiments and collected the data. CL and BB analyzed the data. All authors contributed equally to writing the paper.

## DATA AVAILABILITY STATEMENT

Accession numbers: MT-20 barcodes: GenBank OM858841-OM858852, Transcriptomic samples: BioProject number: PRJNA757218, Growth data: Dryad DOI: https://doi.org/10.5061/dryad.prr4xgxq9

