## Supplemental tables and figures for "Calcium homeostasis disruption initiates rapid growth after micro-fragmentation in the scleractinian coral *Porites lobata*"

**6.1 | Supplementary Figures and Tables**

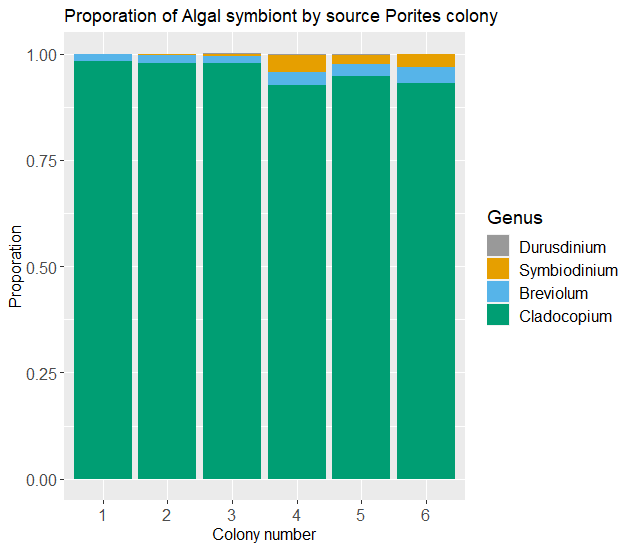

Supplemental Figure 1: Proportion of Symbiodinium Clade by source Porites lobata colony. Proportion of Symbiodinium clades based on highly unique reads (mapping quality > 40) mapping to Clade’s genome.

Supplemental Table 1: Number of sequences in the reference transcriptome after each filtering ste**p**

| Raw reference | rRNAfilter | Long-ORFS | Open reading frames | Alien index | Coral Annotated  (transcripts/genes) | Symbiodinium Annotated (transcripts/gene) |
| --- | --- | --- | --- | --- | --- | --- |
| 1963624 | 1953661 | 811292 | 394555 | 349605 | 196681/55823 | 60475/37404 |

Supplemental Table 2:  The number of raw reads, after quality trim, paired and aligned against reference transcriptome.

| Sample | Raw | After Q>20 | Paired/Alignment rate (%) |
| --- | --- | --- | --- |
| 1t1_S1_R1_001.fastq.gz | 40411887 | 40262269 | 100/96.15 |
| 1t1_S1_R2_001.fastq.gz | 40411887 | 40262269 |  |
| 1t2_S4_R1_001.fastq.gz | 29774025 | 29699824 | 100/94.78 |
| 1t2_S4_R2_001.fastq.gz | 29774025 | 29699824 |  |
| 1t4_S7_R1_001.fastq.gz | 32014830 | 31864908 | 100/93.16 |
| 1t4_S7_R2_001.fastq.gz | 32014830 | 31864908 |  |
| 1t5_S10_R1_001.fastq.gz | 37100965 | 36300314 | 100/92.98 |
| 1t5_S10_R2_001.fastq.gz | 37100965 | 36300314 |  |
| 1t6_S13_R1_001.fastq.gz | 37922178 | 37600580 | 100/97.25 |
| 1t6_S13_R2_001.fastq.gz | 37922178 | 37600580 |  |
| 2t1_S2_R1_001.fastq.gz | 30286530 | 30202343 | 100/95.44 |
| 2t1_S2_R2_001.fastq.gz | 30286530 | 30202343 |  |
| 2t2_S5_R1_001.fastq.gz | 31570608 | 31489229 | 100/97.58 |
| 2t2_S5_R2_001.fastq.gz | 31570608 | 31489229 |  |
| 2t4_S8_R1_001.fastq.gz | 33427925 | 33304499 | 100/95.86 |
| 2t4_S8_R2_001.fastq.gz | 33427925 | 33304499 |  |
| 2t5_S11_R1_001.fastq.gz | 35788659 | 34618172 | 100/94.38 |
| 2t5_S11_R2_001.fastq.gz | 35788659 | 34618172 |  |
| 2t6_S14_R1_001.fastq.gz | 45892227 | 45426834 | 100/96.94 |
| 2t6_S14_R2_001.fastq.gz | 45892227 | 45426834 |  |
| 3t1_S3_R1_001.fastq.gz | 35610993 | 35469639 | 100/93.29 |
| 3t1_S3_R2_001.fastq.gz | 35610993 | 35469639 |  |
| 3t2_S6_R1_001.fastq.gz | 35848204 | 35708680 | 100/94.14 |
| 3t2_S6_R2_001.fastq.gz | 35848204 | 35708680 |  |
| 3t4_S9_R1_001.fastq.gz | 31596203 | 31360005 | 100/93.1 |
| 3t4_S9_R2_001.fastq.gz | 31596203 | 31360005 |  |
| 3t5_S12_R1_001.fastq.gz | 44277047 | 43809930 | 100/95.65 |
| 3t5_S12_R2_001.fastq.gz | 44277047 | 43809930 |  |
| 3t6_S15_R1_001.fastq.gz | 38370519 | 38199360 | 100/96.06 |
| 3t6_S15_R2_001.fastq.gz | 38370519 | 38199360 |  |
| 4t1_S15_R1_001.fastq.gz | 44557694 | 44149932 | 100/96.79 |
| 4t1_S15_R2_001.fastq.gz | 44557694 | 44149932 |  |
| 4t2_S2_R1_001.fastq.gz | 44324757 | 43865923 | 100/95.05 |
| 4t2_S2_R2_001.fastq.gz | 44324757 | 43865923 |  |
| 4t4_S11_R1_001.fastq.gz | 22963920 | 22823537 | 100/97.44 |
| 4t4_S11_R2_001.fastq.gz | 22963920 | 22823537 |  |
| 4t5_S14_R1_001.fastq.gz | 39098341 | 37387406 | 100/95.25 |
| 4t5_S14_R2_001.fastq.gz | 39098341 | 37387406 |  |
| 4t6_S5_R1_001.fastq.gz | 51021407 | 50820124 | 100/95.65 |
| 4t6_S5_R2_001.fastq.gz | 51021407 | 50820124 |  |
| 5t1_S1_R1_001.fastq.gz | 20530864 | 20319138 | 100/78.42 |
| 5t1_S1_R2_001.fastq.gz | 20530864 | 20319138 |  |
| 5t2_S9_R1_001.fastq.gz | 6948235 | 6791713 | 100/95.57 |
| 5t2_S9_R2_001.fastq.gz | 6948235 | 6791713 |  |
| 5t4_S12_R1_001.fastq.gz | 26205810 | 25566136 | 100/89.02 |
| 5t4_S12_R2_001.fastq.gz | 26205810 | 25566136 |  |
| 5t5_S3_R1_001.fastq.gz | 29358571 | 29265742 | 100/95.72 |
| 5t5_S3_R2_001.fastq.gz | 29358571 | 29265742 |  |
| 5t6_S7_R1_001.fastq.gz | 21015779 | 20917704 | 100/94.75 |
| 5t6_S7_R2_001.fastq.gz | 21015779 | 20917704 |  |
| 6t1_S6_R1_001.fastq.gz | 32220307 | 31856476 | 100/92.7 |
| 6t1_S6_R2_001.fastq.gz | 32220307 | 31856476 |  |
| 6t2_S10_R1_001.fastq.gz | 45404846 | 44519689 | 100/89.09 |
| 6t2_S10_R2_001.fastq.gz | 45404846 | 44519689 |  |
| 6t4_S13_R1_001.fastq.gz | 42293636 | 42093497 | 100/97.36 |
| 6t4_S13_R2_001.fastq.gz | 42293636 | 42093497 |  |
| 6t5_S4_R1_001.fastq.gz | 35056090 | 34734875 | 100/95.94 |
| 6t5_S4_R2_001.fastq.gz | 35056090 | 34734875 |  |
| 6t6_S8_R1_001.fastq.gz | 56476217 | 56201663 | 100/94.04 |
| 6t6_S8_R2_001.fastq.gz | 56476217 | 56201663 |  |
| Average | 35,245,642.47 | 34,887,671.4 |  |
| Total | 2,114,738,548 | 2,093,260,282 |  |

Supplemental Table 3: Benchmarking Universal Single-Copy Orthologs (BUSCO) table of transcriptome completeness scores based on metazoan lineage.

| BUSCO Notation | Coral transcriptome | Symbiont transcriptome |
| --- | --- | --- |
| Complete Single-copy | 194 / 20.34% | 38 / 22.22% |
| Complete Duplicated | 694 / 72.75% | 4 / 2.34% |
| Fragmented | 21 / 2.2% | 53 / 30.99% |
| Missing | 45 / 4.72% | 76 / 44.44% |
| Total represented | 93.1% |  |

Supplemental Table 4: Table of significantly differentially expressed genes of interest. List of pairwise comparisons, gene categories, gene names, log2-fold change, uniprot IDs, and significance level of significantly differentially expressed genes which have been previously identified as important in development or stress response. * and ** indicate 0.05 and 0.001 False Discovery Rate (FDR), respectively.

| **Gene Category** | **Secondary Category** | **Comparison** | **Gene name** | **Uniprot ID** | **Log2FC** | **Significance** |
| --- | --- | --- | --- | --- | --- | --- |
| **Oxidative stress** | **Heat shock** | 1V2 | 97 kDa heat shock protein | A0A2B4SKN4_STYPI | 2.02365162 | ** |
|  |  | 1V2 | Heat shock cognate 71 kDa protein | A0A2B4S326_STYPI | 1.63961625 | ** |
|  |  | 2V3 | 97 kDa heat shock protein | A0A2B4SKN4_STYPI | -1.2455299 | * |
|  |  | 1V2 | 10 kDa heat shock protein, mitochondrial (Chaperonin 10) | A7RHS8_NEMVE | 1.53990238 | ** |
|  |  | 1V2 | Heat shock 70 kDa protein 12A | A0A2B4RHP1_STYPI | -0.8376805 | * |
|  |  | 1V2 | Heat shock protein HSP 90-alpha 1 | A0A2B4RG70_STYPI | 1.01276698 | * |
|  |  | 1V2 | Activator of 90 kDa heat shock protein ATPase-like 1 | A0A2B4SBB5_STYPI | 0.92151995 | * |
|  |  | 1V2 | DnaJ homolog subfamily C member 10 (DnaJ homolog subfamily C member 16) | A0A3M6UA46_9CNID | 0.67219806 | * |
|  |  | 1V2 | DnaJ-like subfamily C member 8 | A0A2B4SZE5_STYPI | 0.73690999 | * |
|  |  | 1V2 | DnaJ-like subfamily C member 3 | A0A2B4SDC0_STYPI | 2.51459854 | * |
|  |  | 1V2 | DnaJ-like subfamily A member 2 | A0A2B4SBS3_STYPI | 1.073896 | ** |
|  |  | 4V5 | DnaJ-like subfamily A member 3, mitochondrial | A0A2B4RSU7_STYPI | 0.71052471 | * |
|  |  | 4V5 | DnaJ protein-like 1 (Hsp40) | A0A2B4RDU7_STYPI | 1.48822858 | * |
|  |  | 1V2 | HSP70-1 | D1FX74_CHIFL | 2.95004816 | ** |
| **Oxidative stress** | **Antioxidant** | 1V2 | Thioredoxin | A0A2B4SMK3_STYPI | 1.54949254 | * |
|  |  | 1V2 | Thioredoxin-like_fold domain-containing protein | A0A3M6UVZ5_9CNID | 0.79761591 | * |
|  |  | 1V2 | Thioredoxin, mitochondrial | A0A2B4S014_STYPI | 0.72720729 | * |
|  |  | 1V2 | Peroxiredoxin-1 | A0A2B4S129_STYPI | 1.22216126 | * |
|  |  | 1V2 | Peroxisome proliferator-activated receptor gamma coactivator-related protein 1 | A0A2B4RJR9_STYPI | 1.51120121 | ** |
|  |  | 1V2 | Ferritin (EC 1.16.3.1) | A0A2B4SBC3_STYPI | 1.25384893 | * |
|  |  | 1V2 | Glutathione transferase (EC 2.5.1.18) | A0A2B4SHD3_STYPI | 1.35939527 | ** |
|  |  | 1V2 | Glutathione S-transferase 1 | A0A2B4RJ40_STYPI | 0.91726338 | * |
|  |  | 1V2 | Hypoxia up-regulated protein 1 | A0A2B4SVS9_STYPI | 1.546 | * |
|  |  | 1V5 | Hypoxia up-regulated protein 1 | A0A2B4SVS9_STYPI | 2.068 | ** |
|  |  | 1V2 | Peptide-methionine (S)-S-oxide reductase (EC 1.8.4.11) | A0A3M6U6Z7_9CNID | 1.126 | * |
|  |  | 1V2 | Peptide-methionine (S)-S-oxide reductase (EC 1.8.4.11) | A0A3M6U6Z7_9CNID | 1.806 | ** |
|  |  | 4V5 | Ferredoxin--NADP(+) reductase (EC 1.18.1.6) (NADPH:adrenodoxin oxidoreductase, mitochondrial) | A0A2B4SR94_STYPI | -2.051 | * |
| **Protein** | **Degradation** | 1V2 | RING-type E3 ubiquitin transferase (EC 2.3.2.27) | A0A2B4RLS5_STYPI | 1.153 | ** |
|  |  | 1V2 | Ubiquitin-conjugating enzyme E2 L3 | A0A2B4RQY2_STYPI | 1.233 | ** |
|  |  | 1V2 | PRP19/PSO4 homolog (EC 2.3.2.27) (Pre-mRNA-processing factor 19) (RING-type E3 ubiquitin transferase PRP19) | A0A3M6UJC2_9CNID | 0.914 | ** |
|  |  | 1V2 | Ubiquitin conjugation factor E4 A | A0A2B4RSY9_STYPI | 0.928 | ** |
|  |  | 1V2 | Ubiquitin-fold modifier-conjugating enzyme 1 | A0A2B4STM1_STYPI | 1.123 | ** |
|  |  | 1V2 | Ubiquitin-like domain-containing protein | A0A3M6UAD0_9CNID | 1.123 | * |
|  |  | 1V2 | E3 ubiquitin-protein ligase HERC2 | A0A2B4SA56_STYPI | -1.857 | * |
|  |  | 1V2 | RBR-type E3 ubiquitin transferase (EC 2.3.2.31) | A0A2B4SUB4_STYPI | 1.182 | * |
|  |  | 1V2 | Ubiquitin-like domain-containing protein | A0A3M6UEK6_9CNID | 0.770 | * |
|  |  | 1V2 | HECT-type E3 ubiquitin transferase (EC 2.3.2.26) | A0A2B4RK80_STYPI | 0.825 | * |
|  |  | 1V2 | Ubiquitin-conjugating enzyme E2 variant 2 | T2MCQ1_HYDVU | 0.806 | * |
|  |  | 1V2 | E3 ubiquitin-protein ligase TRIM71 | A0A2B4SE87_STYPI | 0.816 | * |
|  |  | 1V2 | E3 ubiquitin-protein ligase (EC 2.3.2.26) | A0A2B4SI89_STYPI | 0.726 | * |
|  |  | 1V2 | Ubiquitin carboxyl-terminal hydrolase CYLD | A0A2B4RVQ6_STYPI | -1.405 | * |
|  |  | 1V2 | E3 ubiquitin protein ligase (EC 2.3.2.27) | A0A2B4RWL9_STYPI | 0.650 | * |
|  |  | 1V2 | Ubiquitin carboxyl-terminal hydrolase 7 (EC 3.4.19.12) (Ubiquitin thioesterase 7) (Ubiquitin-specific-processing protease 7) | A0A2B4S940_STYPI | 0.800 | * |
|  |  | 1V2 | HECT-type E3 ubiquitin transferase (EC 2.3.2.26) | A0A3M6TI31_9CNID | 0.654 | * |
|  |  | 1V2 | E3 ubiquitin-protein ligase DTX3L | A0A2B4S0G1_STYPI | -1.036 | * |
|  |  | 1V2 | Ubiquitin | A0A2B4SVQ7_STYPI | -0.965 | * |
|  |  | 1V2 | E3 ubiquitin-protein ligase ZSWIM2 | A0A2B4SEP7_STYPI | -0.691 | * |
|  |  | 1V2 | E3 ubiquitin-protein ligase ZNRF2 | A0A2B4STQ2_STYPI | 0.649 | * |
|  |  | 1V2 | Ubiquitin-conjugating enzyme E2 K | A0A2B4T0N3_STYPI | -1.569 | * |
|  |  | 1V2 | 26S proteasome non-ATPase regulatory subunit 6 (26S proteasome regulatory subunit RPN7) | A0A3M6V3Z3_9CNID | 1.271 | * |
|  |  | 1V2 | 26S proteasome regulatory subunit 7 (Proteasome 26S subunit ATPase 2) | A0A3M6UBT2_9CNID | 0.779 | * |
|  |  | 2V3 | E3 ubiquitin-protein ligase TRIM71 | A0A2B4SE87_STYPI | -0.976 | * |
|  |  | 4V5 | Ubiquitin carboxyl-terminal hydrolase MINDY (EC 3.4.19.12) | A0A2B4RUW9_STYPI | 0.855 | * |
|  |  | 4V5 | E3 ubiquitin-protein ligase RNF213 | A0A2B4SMR7_STYPI | -0.977 | * |
|  |  | 4V5 | Ubiquitin-conjugating enzyme E2 variant 2 | T2MCQ1_HYDVU | 0.802 | * |
|  |  | 4V5 | Ubiquitin-conjugating enzyme E2 L3 | A0A2B4RQY2_STYPI | 0.916 | * |
|  |  | 4V5 | Ubiquitin-like protein ATG12 | A0A3M6TTA0_9CNID | 0.782 | * |
|  |  | 4V5 | E3 ubiquitin-protein ligase DTX3L | A0A2B4S0G1_STYPI | -1.162 | * |
|  |  | 4V5 | Ubiquitin carboxyl-terminal hydrolase (EC 3.4.19.12) | A0A3M6TW59_9CNID | 0.701 | * |
|  |  | 4V5 | E3 ubiquitin-protein ligase RNF213 | A0A2B4SK29_STYPI | -0.990 | * |
|  |  | 4V5 | E3 ubiquitin-protein ligase (EC 2.3.2.27) | A0A3M6TW29_9CNID | -1.418 | * |
|  |  | 4V5 | RBR-type E3 ubiquitin transferase (EC 2.3.2.31) | A0A2B4SUB4_STYPI | 1.028 | * |
|  |  | 4V5 | E3 ubiquitin-protein ligase RNF181 | A0A2B4RM06_STYPI | 2.382 | * |
|  |  | 4V5 | Ubiquitin-like domain-containing protein | A0A3M6V0U6_9CNID | 0.934 | * |
| **Protein** | **Synthesis** | 1V2 | Protein disulfide-isomerase TMX3 | A0A2B4SUK4_STYPI | 0.792 | * |
|  |  | 1V2 | Protein disulfide-isomerase (EC 5.3.4.1) | A0A2B4RL32_STYPI | 1.085 | * |
|  |  | 1V2 | Protein disulfide-isomerase (EC 5.3.4.1) | A0A2B4RVE6_STYPI | -1.885 | * |
|  |  | 1V2 | Protein disulfide-isomerase (EC 5.3.4.1) | A0A2B4RUX0_STYPI | 1.923 | * |
|  |  | 1V2 | 5-aminolevulinate synthase (EC 2.3.1.37) (5-aminolevulinic acid synthase) (Delta-ALA synthase) (Delta-aminolevulinate synthase) | A0A3M6V3W3_9CNID | 2.073 | ** |
|  |  | 1V2 | D-aminoacyl-tRNA deacylase (EC 3.1.1.96) | A0A3M6UHT7_9CNID | 1.798 | ** |
|  |  | 1V2 | Lysine--tRNA ligase (EC 6.1.1.6) (Lysyl-tRNA synthetase) | A0A3M6TG26_9CNID | 1.419 | ** |
|  |  | 1V2 | Glutamyl-tRNA synthetase (EC 6.1.1.15) (EC 6.1.1.17) (Prolyl-tRNA synthetase) | A0A2B4RQE5_STYPI | 1.035 | ** |
|  |  | 1V2 | Threonyl-tRNA synthetase (EC 6.1.1.3) | A0A2B4S3J9_STYPI | 1.017 | ** |
|  |  | 1V2 | Leucyl-tRNA synthetase (EC 6.1.1.4) (Fragment) | T2M4D5_HYDVU | 1.087 | ** |
|  |  | 1V2 | Tryptophanyl-tRNA synthetase (EC 6.1.1.2) (Fragment) | A0A3M6UUJ9_9CNID | 0.946 | * |
|  |  | 1V2 | Arginyl-tRNA synthetase (EC 6.1.1.19) | A0A3M6TU20_9CNID | 0.868 | * |
|  |  | 1V2 | Alanine--tRNA ligase (EC 6.1.1.7) | A7SHU9_NEMVE | 1.106 | * |
|  |  | 1V2 | Tyrosine--tRNA ligase, cytoplasmic (EC 6.1.1.1) (Tyrosyl-tRNA synthetase) | A0A2B4SXR4_STYPI | 0.823 | * |
|  |  | 1V2 | Asparagine--tRNA ligase (EC 6.1.1.22) (Fragment) | T2MI28_HYDVU | 0.815 | * |
|  |  | 1V2 | Aminoacyl-tRNA hydrolase (EC 3.1.1.29) | A0A3M6U0G4_9CNID | 1.091 | * |
|  |  | 1V2 | Seryl-tRNA synthetase (EC 6.1.1.11) | A0A3M6UPT9_9CNID | 0.888 | * |
|  |  | 1V2 | Glutaminyl-tRNA synthetase | A0A2B4RHS6_STYPI | 1.112 | * |
|  |  | 1V2 | Cytoplasmic tRNA 2-thiolation protein 2 | A0A3M6UGL7_9CNID | 0.822 | * |
|  |  | 1V2 | Aspartate--tRNA ligase, cytoplasmic (EC 6.1.1.12) (Aspartyl-tRNA synthetase) | A0A3M6TLW8_9CNID | 0.743 | * |
|  |  | 2V3 | Elongation factor Tu | A0A3M6TB99_9CNID | -0.886 | * |
|  |  | 4V5 | Aspartate aminotransferase (EC 2.6.1.1) | A0A3M6UMV2_9CNID | 0.999 | ** |
|  |  | 4V5 | Branched-chain-amino-acid aminotransferase-like protein 2 | A0A2B4SFE9_STYPI | 1.265 | * |
|  |  | 4V5 | 5-aminoimidazole-4-carboxamide ribonucleotide formyltransferase (EC 2.1.2.3) (EC 3.5.4.10) (AICAR transformylase) (AICAR transformylase/inosine monophosphate cyclohydrolase) (Bifunctional purine biosynthesis protein ATIC) (IMP synthase) (Inosinicase) (Phosphoribosylaminoimidazolecarboxamide formyltransferase) | A0A2B4SE86_STYPI | 0.751 | * |
|  |  | 4V5 | Aminotran_1_2 domain-containing protein | A0A3M6THP0_9CNID | 0.661 | * |
|  |  | 4V5 | Tyrosine aminotransferase (TAT) (EC 2.6.1.5) | A0A3M6U7B9_9CNID | -0.951 | * |
|  |  | 4V5 | Aminotran_1_2 domain-containing protein | A0A3M6THP4_9CNID | 0.629 | * |
|  |  | 4V5 | Ribosome production factor 2 homolog (Ribosome biogenesis protein RPF2 homolog) | A0A3M6TEH0_9CNID | 0.853 | * |
|  |  | 4V5 | Ribosomal protein S6 kinase (EC 2.7.11.1) | A0A3M6UDF5_9CNID | 0.996 | * |
|  |  | 4V5 | Ribosomal RNA small subunit methyltransferase NEP1 | A0A2B4SHV4_STYPI | 0.715 | * |
|  |  | 4V5 | D-aminoacyl-tRNA deacylase (EC 3.1.1.96) | A0A3M6UHT7_9CNID | 0.958 | * |
|  |  | 4V5 | Cytoplasmic tRNA 2-thiolation protein 2 | A0A3M6UGL7_9CNID | 0.819 | * |
|  |  | 4V5 | Tryptophanyl-tRNA synthetase (EC 6.1.1.2) (Fragment) | A0A3M6UUJ9_9CNID | 0.700 | * |
|  |  | 4V5 | Aspartate--tRNA ligase, cytoplasmic (EC 6.1.1.12) (Aspartyl-tRNA synthetase) | A0A3M6TLW8_9CNID | 0.687 | * |
|  |  | 4V5 | Lysine--tRNA ligase (EC 6.1.1.6) (Lysyl-tRNA synthetase) | A0A3M6TG26_9CNID | 0.840 | * |
| **Protein** | **Transport** | 1V2 | B(0,+)-type amino acid transporter 1 | A0A2B4RS97_STYPI | 1.175 | * |
|  |  | 1V2 | Putative sodium-coupled neutral amino acid transporter 10 | A0A2B4RSJ0_STYPI | 0.839 | * |
|  |  | 1V2 | High affinity cationic amino acid transporter 1 | A0A2B4SC16_STYPI | 1.016 | ** |
|  |  | 4V5 | 60S ribosomal export protein NMD3 (Fragment) | A7SR71_NEMVE | 0.769 | * |
| **Protein** | **Translation initiation** | 1V2 | Eukaryotic translation initiation factor 3 subunit D (eIF3d) (Eukaryotic translation initiation factor 3 subunit 7) | A0A3M6T603_9CNID | 1.476 | ** |
|  |  | 1V2 | Eukaryotic translation initiation factor 3 subunit C (eIF3c) (Eukaryotic translation initiation factor 3 subunit 8) | A0A2B4RS17_STYPI | 1.246 | ** |
|  |  | 1V2 | Eukaryotic translation initiation factor 3 subunit B (eIF3b) (Eukaryotic translation initiation factor 3 subunit 9) | A0A2B4SKJ2_STYPI | 1.227 | ** |
|  |  | 1V2 | Eukaryotic translation initiation factor 4E | A0A2B4S6J0_STYPI | 1.263 | ** |
|  |  | 1V2 | Eukaryotic translation initiation factor 3 subunit F | A0A2B4SKE2_STYPI | 1.423 | ** |
|  |  | 1V2 | Eukaryotic translation initiation factor 3 subunit F (eIF3f) (Eukaryotic translation initiation factor 3 subunit 5) | A7SIV6_NEMVE | 1.163 | ** |
|  |  | 1V2 | Eukaryotic translation initiation factor 3 subunit L (eIF3l) | A0A2B4RG74_STYPI | 0.985 | ** |
|  |  | 1V2 | Eukaryotic translation initiation factor 3 subunit I (eIF3i) | A0A3M6UR37_9CNID | 0.944 | ** |
|  |  | 1V2 | Eukaryotic translation initiation factor 6 (eIF-6) | A0A2B4SAD2_STYPI | 0.803 | * |
|  |  | 1V2 | Eukaryotic translation initiation factor 3 subunit G (eIF3g) (Eukaryotic translation initiation factor 3 RNA-binding subunit) (eIF-3 RNA-binding subunit) (Eukaryotic translation initiation factor 3 subunit 4) | A0A2B4SEK6_STYPI | 0.943 | * |
|  |  | 1V2 | Eukaryotic translation initiation factor 4 gamma 2 | A0A2B4S5I7_STYPI | 0.776 | * |
|  |  | 1V2 | Eukaryotic translation initiation factor 4B | A0A2B4RI97_STYPI | 0.809 | * |
|  |  | 1V2 | General transcription factor TFIIB (Transcription initiation factor IIB) | A0A2B4SLJ9_STYPI | 0.694 | * |
|  |  | 1V2 | Eukaryotic translation initiation factor 2 subunit 2 | A0A2B4S0K0_STYPI | 0.754 | * |
|  |  | 1V2 | Eukaryotic translation initiation factor 3 subunit E (eIF3e) (Eukaryotic translation initiation factor 3 subunit 6) | A0A2B4RUN6_STYPI | 0.680 | * |
|  |  | 1V2 | Transcription initiation factor TFIID subunit 10 | A0A2B4RQR4_STYPI | 0.742 | * |
|  |  | 1V2 | Eukaryotic translation initiation factor 2D (Ligatin) | A0A3M6TFL1_9CNID | 0.814 | * |
| **Apoptosis** |  | 1V2 | Programmed cell death protein 6 | A0A2B4RNN3_STYPI | 0.649 | * |
|  |  | 1V2 | Caspase-3 | A0A2B4SWS7_STYPI | 0.620 | * |
|  |  | 1V2 | Apoptosis regulator BAX | A0A2B4SFI9_STYPI | 1.081 | * |
|  |  | 1V2 | Bifunctional apoptosis regulator | A0A2B4SJG8_STYPI | 0.849 | * |
|  |  | 4V5 | Caspase-3 | A0A2B4SWS7_STYPI | 0.775 | * |
|  |  | 4V5 | Caspase-7 | A0A2B4S7Z0_STYPI | 1.025 | * |
|  |  | 4V5 | Programmed cell death protein 6 | A0A2B4RNN3_STYPI | 1.048 | ** |
|  |  | 4V5 | Apoptosis regulator BAX | A0A2B4SFI9_STYPI | 0.837 | * |
| **Cell cycle** | **Growth factor** | 1V2 | Epidermal growth factor-like protein 6 | A0A2B4RCT4_STYPI | -1.172 | * |
|  |  | 1V2 | Fibroblast growth factor (FGF) | A0A3M6TGC7_9CNID | -0.793 | * |
|  |  | 1V2 | Fibroblast growth factor receptor 1 | A0A2B4RSX6_STYPI | -1.185 | * |
|  |  | 1V2 | Transforming growth factor 3 protein | A0A0A8K7M1_ACRDI | -0.851 | * |
|  |  | 2V3 | Thyrotroph embryonic factor | A0A2B4S9Y9_STYPI | 2.630 | ** |
|  |  | 2V3 | Prokineticin receptor 1 | A0A2B4RTA0_STYPI | 1.177 | * |
|  |  | 2V3 | Tyrosine-protein kinase transmembrane receptor ROR2 | A0A2B4T378_STYPI | 1.988 | * |
|  |  | 2V3 | Fibroblast growth factor receptor 1-A | A0A2B4RFB7_STYPI | 2.407 | ** |
| **Cytoskeleton** |  | 1V2 | Beta-actin | J7FRQ2_FIMAN | -1.433 | * |
|  |  | 1V2 | Tubulin alpha chain (Fragment) | Q2F6H7_ANTEL | 0.836 | * |
|  |  | 1V2 | Tubulin alpha-1C chain | A0A2B4RTB1_STYPI | 0.839 | * |
|  |  | 1V2 | Alpha-tubulin (Fragment) | Q58HH3_HYDEC | 1.093 | * |
|  |  | 1V2 | Tau-tubulin kinase 1 | A0A2B4RJC1_STYPI | 0.750 | * |
|  |  | 1V2 | Carbonic anhydrase (EC 4.2.1.1) | A0A3M6U8M2_9CNID | -1.732 | ** |
|  |  | 1V2 | Carbonic anhydrase (EC 4.2.1.1) | A0A2B4S8W1_STYPI | -1.798 | * |
|  |  | 1V2 | Integrin alpha-V | A0A2B4RUW4_STYPI | 1.171 | * |
|  |  | 2V3 | SWI/SNF-related matrix-associated actin-dependent regulator of chromatin subfamily A member 5 | A0A2B4SUJ3_STYPI | -0.842 | * |
|  |  | 2V3 | Fibrillar collagen (Fragment) | A1XVT2_HYDVU | -2.979 | * |
|  |  | 2V3 | Matrilin-3 | A0A2B4SV97_STYPI | -1.514 | * |
|  |  | 2V3 | Procollagen-lysine 5-dioxygenase (EC 1.14.11.4) | A0A3M6T4Z7_9CNID | -0.774 | * |
|  |  | 2V3 | Coactosin-like protein | A0A2B4SSA2_STYPI | -1.891 | * |
|  |  | 2V3 | Procollagen-lysine 5-dioxygenase (EC 1.14.11.4) | A0A3M6T4Z7_9CNID | -0.774 | * |
|  |  | 4V5 | Tubulin alpha chain (Fragment) | Q2F6H7_ANTEL | 0.755 | * |
|  |  | 4V5 | Microtubule-actin cross-linking factor 1, isoforms 1/2/3/5 | A0A2B4SNH0_STYPI | -0.640 | * |
| **DNA damage** |  | 4V5 | DNA repair protein | A0A2B4SLG3_STYPI | 1.197 | ** |
|  |  | 4V5 | DNA repair protein RAD51 homolog | A0A2B4SA40_STYPI | 1.278 | * |
|  |  | 2V3 | DNA mismatch repair protein Msh2 | A0A6C0WVN4_ACTEQ | 2.276 | * |
|  |  | 1V2 | DNA topoisomerase I (EC 5.6.2.1) (DNA topoisomerase 1) | A0A2B4SBR9_STYPI | 1.163 | ** |
|  |  | 1V2 | DNA topoisomerase (EC 5.6.2.1) | A0A3M6V2M7_9CNID | 1.532 | ** |
|  |  | 1V2 | DNA double-strand break repair Rad50 ATPase | A0A2B4RUN1_STYPI | -1.062 | * |
| **Calcium** |  | 1V2 | Calcium-transporting ATPase (EC 7.2.2.10) | A0A3M6T8X3_9CNID | 1.517 | ** |
|  |  | 1V2 | Voltage-dependent L-type calcium channel subunit alpha | O97017_STYPI | 0.789 | * |
|  |  | 1V2 | Tyrosine-protein kinase receptor (EC 2.7.10.1) | A0A2B4RSI1_STYPI | 0.808 | * |
|  |  | 1V2 | Calumenin-B | A0A2B4T2P6_STYPI | 1.611 | * |
|  |  | 1V2 | Calreticulin | A0A346HHC6_9CNID | 2.311 | * |
|  |  | 1V2 | Calmodulin | A0A2B4SCT2_STYPI | 1.489 | * |
|  |  | 2V3 | Calmodulin | A0A2B4SCT2_STYPI | -1.396 | * |
| **Calcium** | **Phosphatidylinositol signaling** | 1V2 | Phosphatidylinositol transfer protein beta isoform | A0A2B4SGX6_STYPI | 1.276 | ** |
|  |  | 1V2 | Phosphatidylinositol 3,4,5-trisphosphate 3-phosphatase TPTE2 | A0A2B4SN32_STYPI | 1.612 | ** |
|  |  | 1V2 | Phosphatidylinositol-4,5-bisphosphate 4-phosphatase (EC 3.1.3.78) | A0A2B4S1C9_STYPI | 0.808 | * |
|  |  | 1V2 | Phosphatidylinositol-binding clathrin assembly protein | A0A2B4T1I6_STYPI | 0.768 | * |
|  |  | 1V2 | D-inositol 3-phosphate glycosyltransferase | A0A2B4RXX4_STYPI | 0.778 | * |
|  |  | 1V2 | Diphosphoinositol polyphosphate phosphohydrolase 1 | A0A2B4SXK6_STYPI | 0.894 | * |
|  |  | 1V2 | Phosphatidylinositol 4-kinase type 2 (EC 2.7.1.67) | A0A2B4RZ42_STYPI | 0.724 | * |
|  |  | 1V2 | Type I inositol 1,4,5-trisphosphate 5-phosphatase | A0A2B4RNA5_STYPI | 0.687 | * |
|  |  | 1V2 | Phosphatidylinositol-3-phosphate phosphatase (EC 3.1.3.48) (EC 3.1.3.64) | A0A3M6TG06_9CNID | 0.655 | * |
|  |  | 1V2 | Phosphoinositide phospholipase C (EC 3.1.4.11) | A0A2B4RY04_STYPI | 2.100 | * |
|  |  | 1V2 | Peroxisome proliferator-activated receptor gamma coactivator-related protein 1 | A0A2B4RJR9_STYPI | 1.511 | ** |
|  |  | 1V2 | Serine/threonine-protein kinase receptor (EC 2.7.11.30) | A0A2B4SU78_STYPI | 0.894 | * |
| **Lipid metabolism** |  | 1V2 | Acyl-coenzyme A thioesterase THEM4 | A0A2B4STD7_STYPI | 1.448 | ** |
|  |  | 1V2 | O-acyltransferase | A7RML0_NEMVE | -1.721 | * |
|  |  | 4V5 | Lipid droplet-associated hydrolase (Lipid droplet-associated serine hydrolase) | A0A3M6U228_9CNID | 0.938 | ** |
|  |  | 4V5 | Elongation of very long chain fatty acids protein (EC 2.3.1.199) (Very-long-chain 3-oxoacyl-CoA synthase) | A0A3M6U3H4_9CNID | 1.456 | ** |
|  |  | 4V5 | Sphingolipid 4-desaturase (EC 1.14.19.17) | A0A3M6TES2_9CNID | 1.163 | ** |
|  |  | 4V5 | 3-hydroxyacyl-CoA dehydrogenase type-2 | A0A2B4SYP0_STYPI | 0.930 | * |
| **Carbohydrate metabolism** | **Glycolysis** | 1V2 | Phosphoglycerate mutase (EC 5.4.2.12) (2,3-diphosphoglycerate-independent) | A0A2B4RLF3_STYPI | 1.934 | ** |
|  |  | 2V3 | Phosphoglycerate mutase (2,3-diphosphoglycerate-independent) (EC 5.4.2.12) | A0A2B4RLF3_STYPI | -1.122 | * |
| **Carbohydrate metabolism** | **Citric acid cycle** | 1V2 | Aconitate hydratase, mitochondrial (Aconitase) (EC 4.2.1.-) | A0A3M6UMV8_9CNID | 1.008 | ** |
|  |  | 1V2 | Mitochondrial pyruvate carrier | A0A3M6TNI8_9CNID | 1.111 | ** |
|  |  | 1V2 | Pyruvate dehydrogenase E1 component subunit alpha (EC 1.2.4.1) | A0A3M6UH40_9CNID | 0.704 | * |
|  |  | 1V2 | Dihydrolipoamide acetyltransferase component of pyruvate dehydrogenase complex (EC 2.3.1.-) | A0A2B4SNL5_STYPI | 0.763 | * |
|  |  | 1V2 | Succinate--CoA ligase [GDP-forming] subunit beta, mitochondrial (EC 6.2.1.4) (GTP-specific succinyl-CoA synthetase subunit beta) (G-SCS) (GTPSCS) (Succinyl-CoA synthetase beta-G chain) (SCS-betaG) | A0A3M6V3V3_9CNID | 0.664 | * |
|  |  | 4V5 | Succinate dehydrogenase [ubiquinone] cytochrome b small subunit | A0A3M6V4T9_9CNID | 1.093 | ** |
|  |  | 4V5 | Succinate dehydrogenase (quinone) (EC 1.3.5.1) (Fragment) | A7SCJ3_NEMVE | 1.142 | ** |
|  |  | 4V5 | Malate dehydrogenase (EC 1.1.1.37) | A0A2B4RQH9_STYPI | 0.789 | * |
|  |  | 4V5 | Mitochondrial pyruvate carrier | A0A3M6UPW1_9CNID | 1.618 | * |
|  |  | 4V5 | Mitochondrial pyruvate carrier | A0A3M6TNI8_9CNID | 0.769 | * |
| **Cellular transport** | **Protein transport** | 1V2 | Rab GDP dissociation inhibitor | A0A2B4SJQ6_STYPI | 0.874 | * |
|  |  | 1V2 | Ras-related protein Rab-32A | A0A2B4RQV9_STYPI | -1.317 | * |
|  |  | 1V2 | Rab-3A-interacting protein | A0A2B4SGR3_STYPI | 0.980 | * |
|  |  | 1V2 | Ras-related protein Rab-36 | A0A2B4SF06_STYPI | 0.877 | * |
|  |  | 1V2 | Ras-related protein Rab-2 | A0A2B4SQJ0_STYPI | 0.783 | * |
|  |  | 1V2 | Ras-related protein Rab-7L1 | A0A2B4SHI3_STYPI | 1.020 | * |
| **Cellular transport** | **Proton transport** | 1V2 | Vacuolar proton pump subunit B (V-ATPase subunit B) (Vacuolar proton pump subunit B) | A0A2B4S736_STYPI | 1.045 | ** |
|  |  | 1V2 | V-type proton ATPase 21 kDa proteolipid subunit | A0A2B4SQV4_STYPI | 0.941 | * |
|  |  | 1V2 | Proton-translocating NAD(P)(+) transhydrogenase (EC 7.1.1.1) | A0A2B4SX48_STYPI | 0.739 | * |
|  |  | 1V2 | V-type proton ATPase subunit E | A0A2B4SI00_STYPI | 0.727 | * |
|  |  | 4V5 | V-type proton ATPase 21 kDa proteolipid subunit | A0A2B4SQV4_STYPI | 0.880 | * |
|  |  | 4V5 | V-type proton ATPase subunit C | A7RL79_NEMVE | 0.656 | * |
| **Cellular transport** |  | 1V2 | Sodium/potassium-transporting ATPase subunit beta-1-interacting protein 1 | A0A2B4SDN7_STYPI | 0.939 | * |
|  |  | 1V2 | Transportin-1 | A0A2B4SM89_STYPI | 1.381 | * |
|  |  | 1V2 | Magnesium transporter protein 1 | A0A2B4T0L4_STYPI | 0.860 | * |
|  |  | 1V2 | Major facilitator superfamily domain-containing protein 5 (Molybdate transporter 2 homolog) (Molybdate-anion transporter) (Fragment) | A7SG46_NEMVE | -0.778 | * |
|  |  | 1V2 | Copper transporter | A0A2B4S5R8_STYPI | 1.685 | * |
|  |  | 1V2 | Solute carrier organic anion transporter family member | A0A3M6TG97_9CNID | 0.997 | * |
|  |  | 1V2 | Solute carrier organic anion transporter family member | A0A2B4S483_STYPI | 0.630 | * |
|  |  | 1V2 | Phospholipid-transporting ATPase (EC 7.6.2.1) | A0A2B4S7M6_STYPI | 0.643 | * |
|  |  | 1V2 | Glycine betaine transporter OpuD | A0A2B4RX87_STYPI | 1.088 | * |
|  |  | 1V2 | Protein transport protein SEC23 | A0A3M6U474_9CNID | 0.638 | * |
|  |  | 1V2 | Putative sodium-coupled neutral amino acid transporter 10 | A0A2B4RSJ0_STYPI | 0.839 | * |
|  |  | 1V2 | Solute carrier family 30 member 9 (Zinc transporter 9) | A0A3M6TT97_9CNID | 0.775 | * |
|  |  | 1V2 | Solute carrier family 30 member 9 (Zinc transporter 9) | A0A3M6TT97_9CNID | 0.775 | * |
|  |  | 1V2 | Zinc transporter ZIP11 | A0A2B4SI40_STYPI | 0.688 | * |
|  |  | 1V2 | Zinc transporter ZIP13 | A0A2B4RI41_STYPI | 0.863 | * |
|  |  | 1V2 | Zinc transporter SLC39A7 | A0A2B4RYI4_STYPI | 2.294 | * |
|  |  | 1V3 | Vesicle-fusing ATPase (EC 3.6.4.6) | A7SJ61_NEMVE | 1.018 | ** |
|  |  | 2V3 | Monocarboxylate transporter 10 | A0A2B4S850_STYPI | -1.079 | * |
|  |  | 4V5 | Zinc transporter ZIP11 | A0A2B4SI40_STYPI | 0.734 | * |
|  |  | 4V5 | Zinc transporter 2 | A0A2B4RP90_STYPI | 1.189 | ** |
|  |  | 4V5 | Sodium/potassium-transporting ATPase subunit beta-1 | A0A2B4SP72_STYPI | 1.167 | ** |
|  |  | 4V5 | ABC transporter domain-containing protein (Fragment) | A0A3M6TFU8_9CNID | 1.255 | * |
|  |  | 4V5 | Mitochondrial coenzyme A transporter SLC25A42 | A0A2B4RV20_STYPI | 0.800 | * |
|  |  | 4V5 | Sodium/potassium-transporting ATPase subunit alpha (Fragment) | A0A3M6TW19_9CNID | 1.038 | * |
|  |  | 4V5 | Sodium-and chloride-dependent GABA transporter 2 | A0A2B4S0L0_STYPI | 0.963 | * |
|  |  | 4V5 | Monocarboxylate transporter 10 | A0A2B4RV02_STYPI | 0.736 | * |
|  |  | 4V5 | Sodium-coupled monocarboxylate transporter 1 | A0A2B4SZY2_STYPI | 0.778 | * |
| **RNA/DNA** | **Transcription** | 1V2 | RNA helicase (EC 3.6.4.13) | T2M5C3_HYDVU | 1.629 | ** |
|  |  | 1V2 | RNA helicase (EC 3.6.4.13) | A0A2B4RYD4_STYPI | -1.431 | ** |
|  |  | 1V2 | RNA helicase (EC 3.6.4.13) | A0A3M6TB93_9CNID | 2.679 | ** |
|  |  | 1V2 | RNA helicase (EC 3.6.4.13) | A0A3M6UH79_9CNID | 1.019 | ** |
|  |  | 1V2 | RNA helicase (EC 3.6.4.13) | A0A3M6TE03_9CNID | 1.008 | ** |
|  |  | 1V2 | RNA helicase (EC 3.6.4.13) | J3T9N7_FIMAN | 0.935 | ** |
|  |  | 1V2 | RNA helicase (EC 3.6.4.13) | A0A2B4SNV4_STYPI | 0.910 | ** |
|  |  | 1V2 | RNA helicase (EC 3.6.4.13) | A0A2B4RCN6_STYPI | -1.190 | * |
|  |  | 1V2 | RNA helicase (EC 3.6.4.13) | A0A3M6U918_9CNID | 1.985 | * |
|  |  | 1V2 | RNA helicase (EC 3.6.4.13) | A0A2B4R8X2_STYPI | 0.981 | * |
|  |  | 1V2 | RNA helicase (EC 3.6.4.13) | A0A2B4SVS7_STYPI | 0.769 | * |
|  |  | 1V2 | RNA helicase (EC 3.6.4.13) | A0A3M6U6L2_9CNID | -0.774 | * |
|  |  | 4V5 | RNA helicase (EC 3.6.4.13) | T2MDB0_HYDVU | -1.254 | * |
|  |  | 4V5 | RNA-directed RNA polymerase (EC 2.7.7.48) | A0A2B4RG40_STYPI | -1.565 | * |
|  |  | 4V5 | RNA-directed DNA polymerase from mobile element jockey | A0A2B4SBK8_STYPI | -1.086 | * |
|  |  | 4V5 | RNA-directed DNA polymerase from mobile element jockey | A0A2B4S1E6_STYPI | -1.212 | * |
|  |  | 4V5 | RNA-directed DNA polymerase from mobile element jockey | A0A2B4T008_STYPI | -1.327 | * |
|  |  | 4V5 | RNA-directed DNA polymerase (EC 2.7.7.49) | A0A2B4SZG6_STYPI | -0.889 | * |
| **RNA/DNA** | **Synthesis** | 4V5 | AIR carboxylase (EC 4.1.1.21) (EC 6.3.2.6) (Phosphoribosylaminoimidazole carboxylase) (Phosphoribosylaminoimidazole-succinocarboxamide synthase) (SAICAR synthetase) | A0A2B4RQJ5_STYPI | 2.521 | ** |
| **Cellular energy** |  | 1V2 | Adenylate kinase (EC 2.7.4.3) (ATP-AMP transphosphorylase) (ATP:AMP phosphotransferase) (Adenylate kinase cytosolic and mitochondrial) (Adenylate monophosphate kinase) | A0A3M6TCQ4_9CNID | 0.932 | ** |
| **Cellular energy** | **ATP generation** | 1V2 | ATP synthase subunit alpha | A0A2B4T1C4_STYPI | 1.136 | ** |
|  |  | 1V2 | ATP synthase subunit gamma | A7RNI6_NEMVE | 0.890 | * |
|  |  | 1V2 | ATP synthase F(0) complex subunit B1, mitochondrial | A0A2B4SSQ3_STYPI | 1.210 | * |
|  |  | 1V2 | ATP synthase F(0) complex subunit B1, mitochondrial | A0A2B4SSQ3_STYPI | 1.210 | * |
|  |  | 1V2 | ATP synthase subunit beta (EC 7.1.2.2) | A0A2B4SJB0_STYPI | 0.897 | * |
|  |  | 4V5 | ATP synthase subunit alpha | A0A2B4T1C4_STYPI | 0.839 | * |
| **Cellular energy** | **Electron transport chain** | 1V2 | Electron transfer flavoprotein subunit alpha (Alpha-ETF) | A0A3M6U8Q7_9CNID | 0.935 | * |
|  |  | 1V2 | Complex I-9kD (NADH dehydrogenase [ubiquinone] flavoprotein 3, mitochondrial) (NADH-ubiquinone oxidoreductase 9 kDa subunit) | A0A2B4SMJ8_STYPI | 0.834 | * |
|  |  | 1V2 | Complex I-30kD (NADH dehydrogenase [ubiquinone] iron-sulfur protein 3, mitochondrial) (NADH-ubiquinone oxidoreductase 30 kDa subunit) | A0A3M6TYS4_9CNID | 0.731 | * |
|  |  | 1V2 | NADH dehydrogenase [ubiquinone] flavoprotein 2, mitochondrial | A0A2B4SFU3_STYPI | 0.697 | * |
|  |  | 1V2 | Complex I-B14.7 (NADH dehydrogenase [ubiquinone] 1 alpha subcomplex subunit 11) (NADH-ubiquinone oxidoreductase subunit B14.7) | A0A3M6TFB3_9CNID | 0.686 | * |
|  |  | 1V2 | Complex I-19kD (NADH dehydrogenase [ubiquinone] 1 alpha subcomplex subunit 8) (NADH-ubiquinone oxidoreductase 19 kDa subunit) | A0A3M6TW27_9CNID | 0.672 | * |
|  |  | 1V2 | Complex I-ESSS (NADH dehydrogenase [ubiquinone] 1 beta subcomplex subunit 11, mitochondrial) (NADH-ubiquinone oxidoreductase ESSS subunit) | A0A2B4RK25_STYPI | 0.706 | * |
|  |  | 1V2 | NADH dehydrogenase [ubiquinone] flavoprotein 1, mitochondrial (EC 7.1.1.2) | A0A2B4REL4_STYPI | 0.693 | * |
|  |  | 1V2 | NADH dehydrogenase [ubiquinone] 1 alpha subcomplex subunit 9, mitochondrial | A0A2B4T2D6_STYPI | 0.701 | * |
|  |  | 1V2 | Complex I-49kD (NADH-ubiquinone oxidoreductase 49 kDa subunit) | A0A3M6TNT9_9CNID | 0.800 | * |
|  |  | 1V2 | Complex I-49kD (NADH-ubiquinone oxidoreductase 49 kDa subunit) | A0A2B4S4S7_STYPI | 0.878 | * |
|  |  | 1V2 | Cytochrome c domain-containing protein | A0A3M6UD76_9CNID | 1.462 | ** |
|  |  | 1V2 | Cytochrome c oxidase subunit 5B, mitochondrial | A0A2B4RPW9_STYPI | 1.383 | ** |
|  |  | 1V2 | Cytochrome c oxidase polypeptide VIIc | A0A3M6U6A2_9CNID | 2.770 | * |
|  |  | 1V2 | Cytochrome c oxidase subunit (Cytochrome c oxidase polypeptide VIa) | A0A3M6TGK2_9CNID | 0.984 | * |
|  |  | 1V2 | Cytochrome c oxidase polypeptide Va (Cytochrome c oxidase subunit 5A, mitochondrial) | A0A2B4RZU5_STYPI | 1.172 | * |
|  |  | 1V2 | Cytochrome b(558) alpha chain (Cytochrome b-245 light chain) (Cytochrome b558 subunit alpha) (Neutrophil cytochrome b 22 kDa polypeptide) (Superoxide-generating NADPH oxidase light chain subunit) (p22 phagocyte B-cytochrome) (p22-phox) | A0A3M6TPG0_9CNID | 0.984 | * |
|  |  | 1V2 | Cytochrome b-245 heavy chain | A0A2B4REC7_STYPI | 1.836 | ** |
|  |  | 1V2 | Complex III subunit 8 (Complex III subunit VIII) (Cytochrome b-c1 complex subunit 8) (Ubiquinol-cytochrome c reductase complex 9.5 kDa protein) (Ubiquinol-cytochrome c reductase complex ubiquinone-binding protein QP-C) | A0A3M6UGP1_9CNID | 1.361 | ** |
|  |  | 1V2 | Cytochrome b-c1 complex subunit 6, mitochondrial | A0A2B4SXR9_STYPI | 1.288 | * |
|  |  | 1V2 | Cytochrome b-c1 complex subunit 7 | A0A3M6TEI6_9CNID | 1.917 | * |
|  |  | 1V2 | Cytochrome b5 heme-binding domain-containing protein | A0A3M6U6V7_9CNID | 1.443 | * |
|  |  | 1V2 | Cytochrome b5 heme-binding domain-containing protein (Fragment) | A0A3M6V3E2_9CNID | 0.778 | * |
|  |  | 1V2 | Cytochrome b5 heme-binding domain-containing protein | A0A3M6TUM8_9CNID | 1.102 | * |
|  |  | 1V2 | Cytochrome P450 4F5 | A0A2B4RGC5_STYPI | -0.925 | * |
|  |  | 1V2 | Cytochrome P450 1A1 | A0A2B4S3Z7_STYPI | -0.740 | * |
|  |  | 1V2 | NADPH--cytochrome P450 reductase | A0A2B4S487_STYPI | 0.755 | * |
|  |  | 4V5 | Complex I-49kD (NADH-ubiquinone oxidoreductase 49 kDa subunit) | A0A2B4S4S7_STYPI | 1.503 | ** |
|  |  | 4V5 | Electron transfer flavoprotein-ubiquinone oxidoreductase (ETF-QO) (EC 1.5.5.1) | A0A2B4SE12_STYPI | 1.547 | ** |
|  |  | 4V5 | Cytochrome b-561 (Cytochrome b561) | A0A3M6UV69_9CNID | 1.030 | ** |
| **Other** | **Histone** | 1V2 | Histone deacetylase (EC 3.5.1.98) | A0A2B4SH89_STYPI | 1.470 | ** |
|  |  | 1V2 | Histone H1-delta | A0A2B4SSY5_STYPI | -1.461 | * |
|  |  | 1V2 | Histone-binding protein RBBP7 | A0A2B4SQP1_STYPI | 0.846 | * |
|  |  | 1V2 | Putative histone-lysine N-methyltransferase PRDM6 | A0A2B4SW04_STYPI | 2.729 | * |
|  |  | 1V2 | Histone chaperone asf1 | A0A2B4SNY5_STYPI | 0.776 | * |
|  |  | 1V2 | Histone acetyltransferase (EC 2.3.1.48) | A0A3M6TT41_9CNID | 1.390 | * |
|  |  | 1V2 | Histone acetyltransferase (EC 2.3.1.48) | A0A2B4SF80_STYPI | 0.682 | * |
|  |  | 1V2 | [Histone H3]-trimethyl-L-lysine(4) demethylase (EC 1.14.11.67) | A0A3M6UK63_9CNID | 0.623 | * |
| **Other** | **Immune response** | 1V2 | Complement C3 | A0A2B4SP95_STYPI | -3.003 | * |
|  |  | 1V2 | Complement C1q and tumor necrosis factor-related protein 9B | A0A2B4SXX5_STYPI | -0.775 | * |
